# Persistent Low-Level Infections of Elephant Endotheliotropic Herpesvirus and Elephant Gammaherpesvirus Detected in Skin Nodules and Saliva from Wild and Zoo African Elephants

**DOI:** 10.1101/2024.11.17.624021

**Authors:** Virginia Riddle Pearson, Gary S. Hayward

## Abstract

This novel study detected persistent low-level infection of Elephant Endotheliotropic Herpesviruses (EEHV), that can cause highly pathogenic Elephant Hemorrhagic Disease (EHD) in *Loxodonta* and *Elephas*, and co-infection of presumed less pathogenic Elephant Gammaherpesviruses (EGHV), in skin nodule biopsies, saliva and tissues collected from 43 wild *L. africana* (savannah elephant) in Botswana, Kenya, South Africa and Zimbabwe; in saliva from 25 wild *L. cyclotis* (forest elephant) in Gabon and 7 wild-born *L.africana* at Six Flags Safari Park, USA, over an extended period of seven years; and in saliva, blood and tissues from an additional 200 *L. africana* and 100 *Elephas maximus* in USA zoos. DNA from these samples was extracted in our USA laboratories and amplified by conventional polymerase chain reaction using three-round nested primer sets designed specifically to screen for known EEHV and EGHV genes loci and to discover new species and subtypes. Sanger sequencing of purified DNA from nearly all samples yielded unambiguous positive genetic matches to previously known *Loxodonta*-associated EEHV2, EEHV3A, EEHV3B, EEHV6, EEHV7A, and EGHV1B, EGHV2, EGHV3B, EGHV4B, EGHV5B and discovered novel types EEHV3C-H and EEHV7B and the prototype EGHV1B. Many of the primer sets used could also have detected known *Elephas-*associated EEHV1A, EEHV1B, EEHV4, and EEHV5 if present, but they did not. This extensive EEHV and EGHV sequence library will be a significant contribution to the elephant virology community.

## INTRODUCTION

The last two living Genera of the ancient Order Proboscidea: *Loxodonta* (African elephant) and *Elephas* (Asian elephant) are listed by the IUCN as endangered or critically endangered [1,2,]. Habitat fragmentation, increasing human-elephant conflict, and poaching of elephants for ivory, skin and internal organs have decimated, even eliminated, entire populations in all traditional range countries [3–10]. *Loxodonta* and *Elephas* are further threatened by often fatal Elephant Hemorrhagic Disease (EHD) attributed to Elephant Endotheliotropic Herpesviruses (EEHV) Death can occur in as little as 12 hours after the first appearance of clinical signs such as high-level viremia, cyanosis of the tongue, edema of the head and limbs, lethargy, and tachycardia. Hundreds of confirmed cases of severe or lethal EHD attributed to *Elephas*-associated EEHV1A, EEHV1B, EEHV4 and EEHV5 have been documented in wild and zoo *E.maximus*, including critically endangered sub-species *E.m borneensis* and *E.m sumantranus* [11–34]. More recently, almost two dozen cases of EHD attributed to *Loxodonta*-associated EEHV2, EEHV3A, EEHV3B, and EEHV6 have been documented in *L. africana* in zoos worldwide, but none as yet in range countries [35–42]. Previously, we reported finding evidence of these same four types of *Loxodonta*-associated EEHV in lung biopsies from a wild *L.africana* in Kenya. [43–47] collected for this study. As the numbers of free-ranging *Loxodonat,* continue to decrease precipitously, and herds become increasingly isolated, lethal EHD may become a much more significant threat in wild *Loxodonta* [48].

## THIS STUDY

We commenced this study to look for any evidence of cross-species infection between *Loxodonta* and *Elephas*. The unambiguous distinct types of EEHV we have identified in *L. africana and L. cyclotis* dispel the originally held hypothesis that the severe pathogenesis and mortality attributable to EEHV in *Elephas* might be caused by lethal cross-species infections from *Loxodonta*. EEHV species are an ancient cladal group that evolved along with the ancestors of modern proboscidean hosts for tens of millions of years. They form two major branches with either AT-rich versus GC-rich character, which in turn split into seven distinctive species that diverge overall from each other by at least 16 to 20% at the nucleotide level. Each species also displays A plus B subtypes that may contain up to ten localized non-adjacent chimeric domains (CD) or hypervariable gene blocks encompassing up to seven distinct subtype clusters with large (15 to 50%) protein level differences [49–53]. The five EGHV species group POL genes are widely diverged displaying between 37 and 45% nucleotide polymorphisms and fall into three ancient deep lineages within the mammalian gamma herpesviruses corresponding to estimated last common ancestors (LCA) occurring about 200 million years ago [54–57]. Four of the EGHV species (EGHV1, EGHV3, EGHV4 and EGHV5) further divide into consistent A and B diasporas that differ by around 3 to 5% (15 to 24/450-bp polymorphisms) and are largely specific for *Elephas* versus *Loxodonta* hosts respectively [58,59]. (**Table 1**.)

**Table 1.** EEHV and EGHV Species and Subtypes.

| Species | Case# | Source | Host | Discovery/Subsequent Reports |
| --- | --- | --- | --- | --- |
| *EEHV1A | NAP#11,19 | Bl/Nec, TW | EM | Richman1999; Stanton2009 |
| *EEHV1B | NAP#14,18 | Bl/Nec, TW | EM | Ehlers2001; Fickel2001 |
| *EEHV2 | NAP#12, #42 | Nec, LungN, SkinN | LA | Richman1999; Zong2010; **Pearson2011, 2023** |
| *EEHV3A | NAP#27, #42 | Bl/Nec, LN, SkinN | EM, LA | Garner2007; Zong2010; **Pearson2011**, Fayette2017 |
| **EEHV3B** | **NAP#63** | **Skin Nod, Bl, TW** | **LA** | **Pearson2012**; Bronson, 2013; Fayette 2017 |
| *EEHV4A | NAP#22 | Bl/Nec | EM | Garner2007 |
| EEHV4B | NAP#69 | Bl, TW | EM | Feury2014 |
| *EEHV5A | NAP#29 | Bl, TW | EM | Latimer2008; Atkins2011 |
| EEHV5B | NAP#58 | Bl, TW | EM | Atkins2011 |
| *EEHV6A | NAP#35, #42 | Bl, LungNod, Nec | LA | Latimer2009; Zong2010; **Pearson2011, 2024** |
| EEHV6B |  |  | LA | **Pearson2014** |
| EEHV7A | NAP#42 | LungN, SkinN. | LA | Zong2010; **Pearson2011** |
| **EEHV7B** |  | **Skin N, Sal** | **LA** | **Pearson2011, 2017** |
| **Gammaherpesvirus Sub-Family:** |  |  |  |  |
| EGHV1A |  | Con/Swab, Saliva | EM | Wellehan2007; Ehlers 2007 |
| **EGHV1B** | **NAG#11,12** | **Saliva** | **LA** | **Pearson2012** |
| EGHV2 | NAG#1,13,14 | Con/Swab, Bl | EM, LA | Wellehan2008; Latimer; **Pearson2012** |
| EGHV3A | NAG#2,7 | Con/Swab, Bl | EM | Wellehan2007; Latimer2008 |
| EGHV3B | NAG#5,15,16 | Bl, Sal, Nec | LA | Latimer2008, **Pearson2012, 2017** |
|  | NAG 16, NAG31-32 |  |  |  |
| EGHV4A | NAG58 | Spleen | EM | **Pearson2023** |
| EGHV4B | NAG#17 | Con/Swab, Saliva | LA | Wellehan2007; **Pearson2012** |
| EGHV5A | NAG#6 | Trk Papilloma | EM | Latimer2008; Masters2011 |
| EGHV5B | NAG#8 | Trk Pap, Nec, Sal, N | LA | Long2011; **Pearson2013, 2017, 2024** |
| * = Lethal Acute Hemorrhagic Disease; N = Skin Nodule; Sal = Saliva; TW = Trunk Wash; TrkPap = Trunk papilloma interior; Con = eye conjunctiva swab; Nev = Necropsy tissues and organs |  |  |  |  |
| EM = *Elephas maximus*; LA = *Loxodonta africana* |  |  |  |  |

### Search for evidence of EEHV and EGHV in Skin Nodules and Saliva from wild *L.africana* in Kenya, Botswana, South Africa and Zimbabwe

When we began this study, little was known about herpesviruses in wild *Loxodonta* except for the Jacobson et al 1986 descriptions of viral nuclear inclusion bodies morphologically consistent with herpesviruses in raised cutaneous fibropapillomas (skin nodules) in a group of ninety-nine *L. africana* imported to USA from Zimbabwe in 1982-84, and the McCully et al 1971 findings of white lymphoid nodules containing Cowdry Type A intranuclear inclusions pathognomic of herpesvirus disease in lungs of culled *L.africana* in South Africa [60,61]. We hypothesized that we might find evidence of since-characterized EEHV and EGHV in similar skin nodules if they existed on living wild *L.africana* and *L. cyclotis,* to discover unique genetic polymorphisms that could help understand more clearly the origins and pathology of EHD in *Loxodonta* and *Elephas*. As we had been unable to locate any USA zoo elephants with skin nodules per Jacobson’s descriptions, including still-living members of the 1982-84 importation from Zimbabwe, so in 2009, Pearson began to search for skin nodules on living wild *L.africana* in Kenya, which were found and photographed. (**Figure 1**.) Nodules were visually distinguished from warts based on gross morphology: nodules were round, ulcerative or thinly crusted over, circumscribed, predominantly on the trunk and often of a different appearance than surrounding skin.

**Figure 1.**
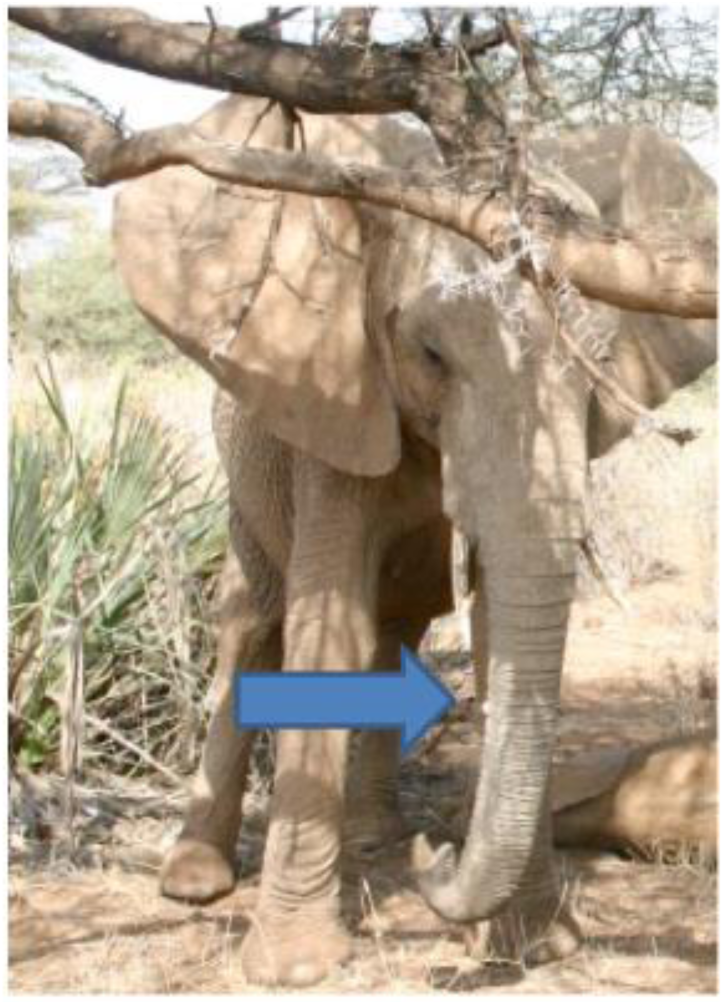
skin nodules on wild *L. Africana,* 2009

In 2011, Pearson returned to Kenya in collaboration with Save The Elephants (STE) in Kenya and Kenya Wildlife Service Veterinary and Capture Service, to search for, immobilize and collect biopsies of skin nodules from three young calves and two subadult wild *L.africana*, albeit, not the same individuals as photographed in 2009. (**Figure 2**, **Figure 3**, **Figure 4**.) Wild elephants are constantly moving and it was very difficult to spot nodules at a distance with binoculars and keep the particular elephant in sight while we approached by vehicle with the tranquilizer. Elephants can be immobilized for only a short duration (preferably less than 25 minutes) and recovery time after antidote is injected is approximately 90 seconds posing imminent danger to the veterinarians and researchers. In order to give us time to collect nodule biopsies from the calves, we immobilized the mothers of the youngest calves first. As such, we did not have sufficient time to collect skin adjacent to the nodules for investigation of EEHV and EGHV cellular tropism [62]. No injuries to elephants or personnel occurred during these field operations.

**Figure 2.**
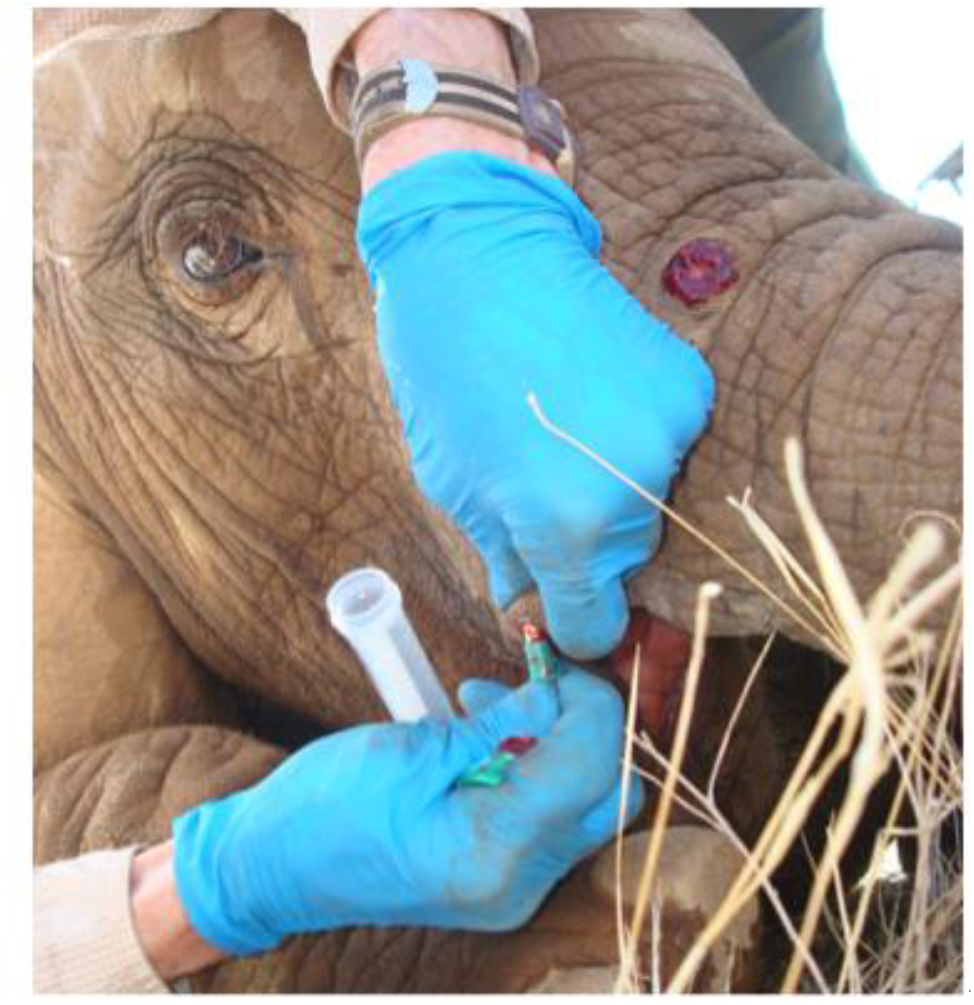
Skin nodule on young calf cGM, Samburu National Reserve, Kenya, 2011. Credit Pearson, 2009 and 2011.

**Figure 3.**
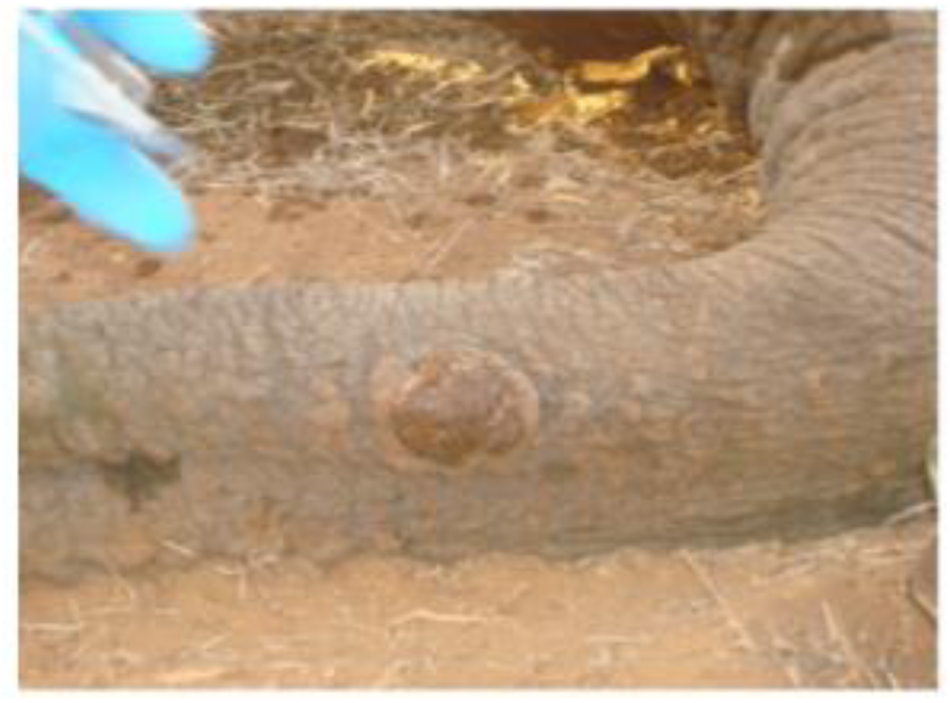
skin nodule on calf cMT.

**Figure 4.**
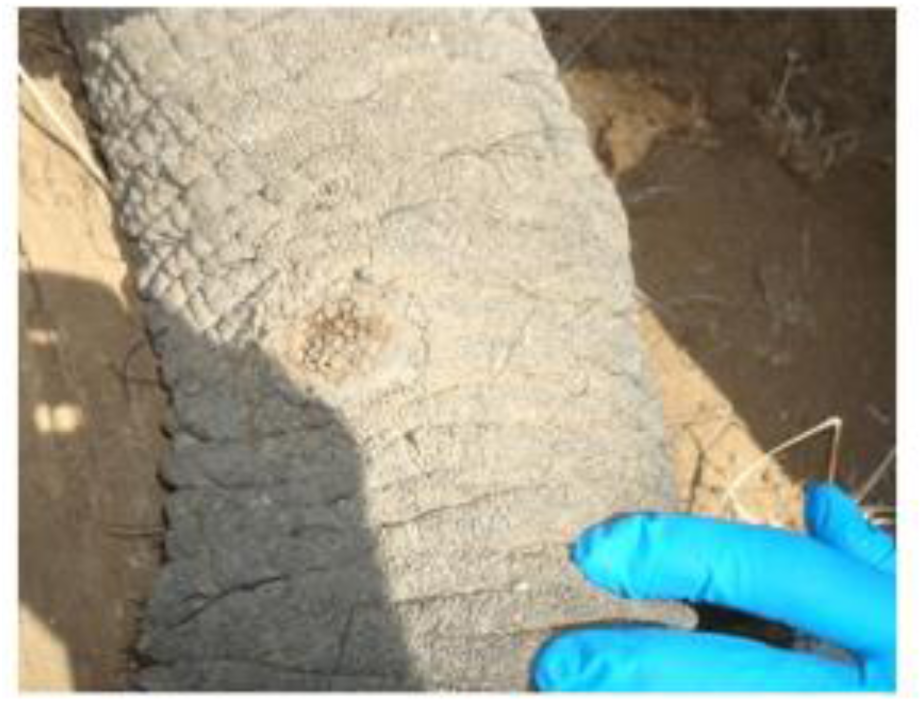
skin nodule on juvenile HIMA. Samburu National Reserve, Kenya. Credit Pearson, 2011.

Pearson also collected the lung tissue biopsies reported in Zong et al 2015 (Genbank accession #KT832496-KT832512) from the sub-adult named Enthusiasm who was poached for her ivory tusks. Enthusiasm was well known to Save the Elephants as a member of the Virtues Family whom they had studied for many years. (**Figure 5**, **Figure 6**, **Figure 7**.) We expected that we might find evidence of since-characterized EEHV and EGHV in this lung, which we did by genetic analysis back in our USA laboratories. [44–47].

**Figure 5.**
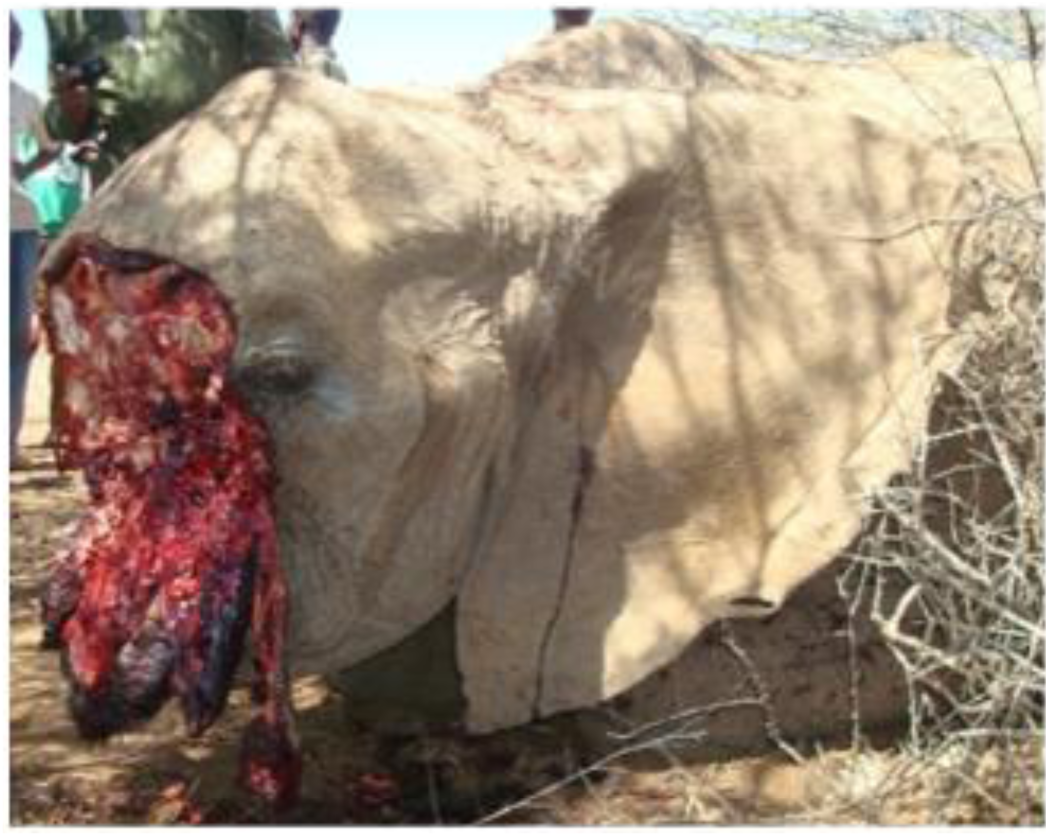
*L. africana* Enthusiasm, Samburu, Kenya.

**Figure 6.**
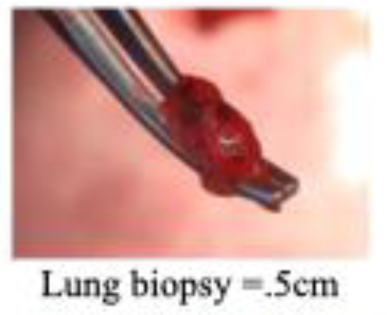
Lung was sectioned and biopsies collected by Pearson within a few hours post-mortem.

**Figure 7.**
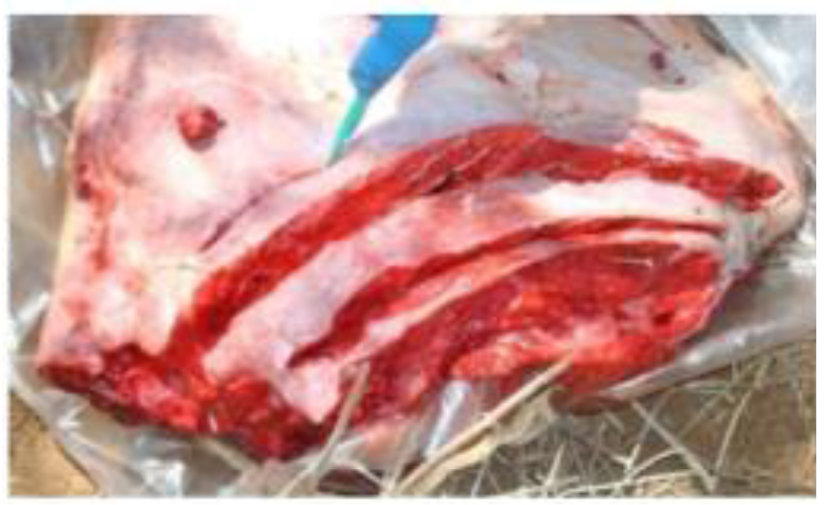
.5 cm lung cubes were immersed in Qiagen RNAlater. Credit Pearson, 2011.

On a second expedition in 2013, in collaboration with Elephants Without Borders (EWB) in Botswana and Zimbabwe, Pearson searched for, immobilized and collected biopsies of skin nodules from two young calves, one of which had four nodules on the underside side of his trunk [44–47]. (**Figure 8**, **Figure 9**)

**Figure 8.**
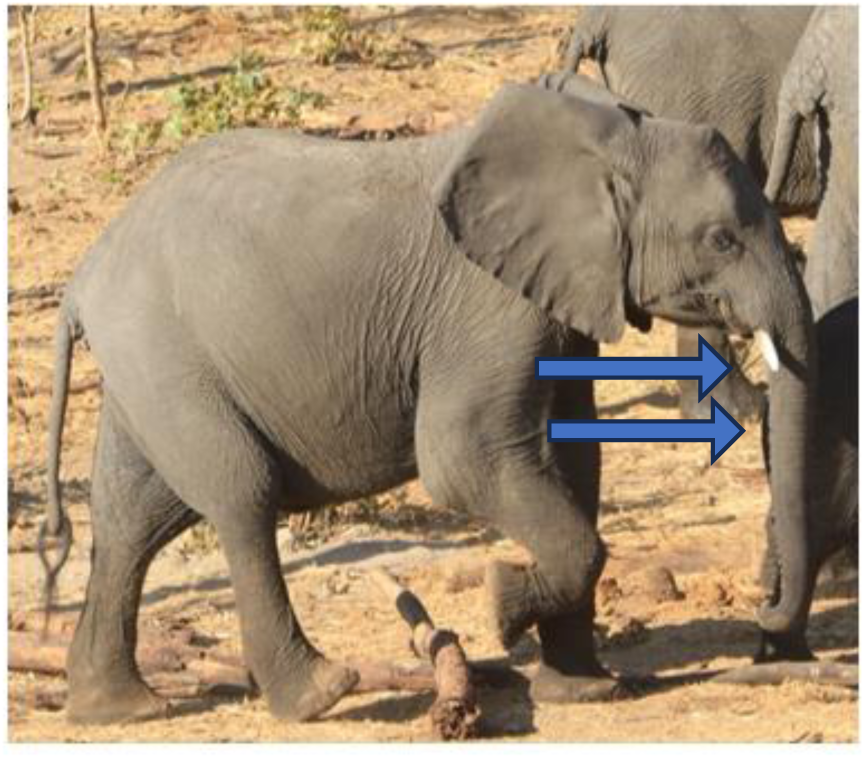
left: Location of four nodules on trunk underside of wild juvenile male *L.africana*, designated BW1M Nod1 to Nod4. Chobe National Park, Kasane, Botswana, 2013.

**Figure 9.**
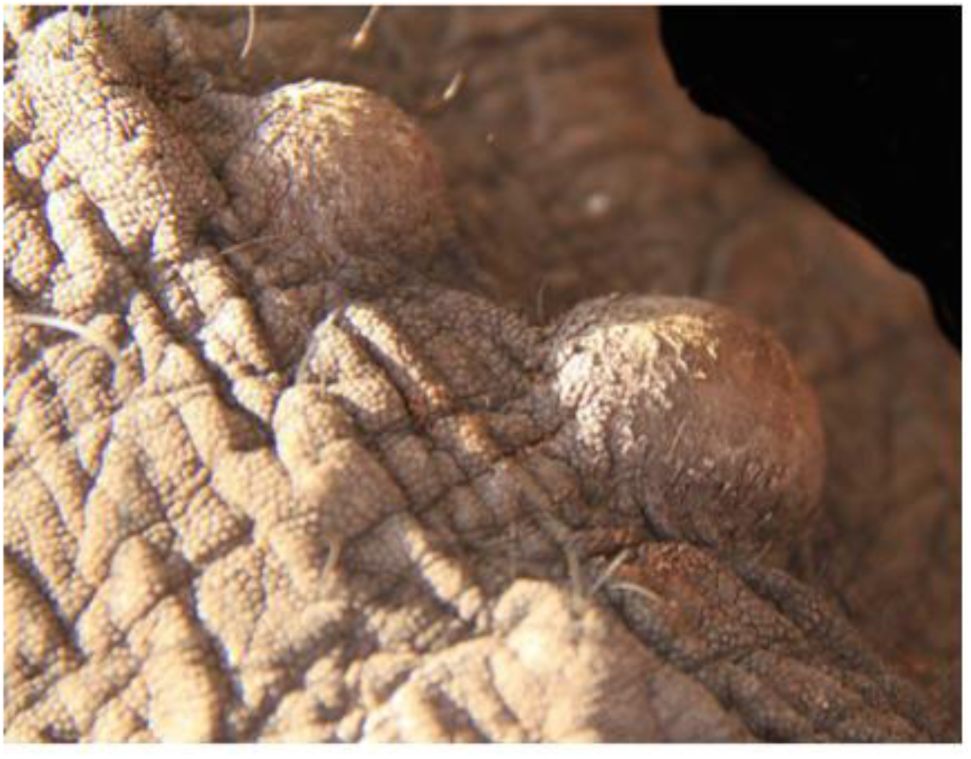
right: 6 mm punch biopsies of BW1M Nod1 were collected by Pearson. Credit Pearson, 2013.

During these two expeditions and another in South Africa with Elephants Alive in 2013, Pearson also collected saliva and non-nodular skin warts (**Figure 13**) from twenty-six *L.africana* who had been immobilized for emergency treatment of gunshot wounds or GPS collaring purposes or who were semi-habituated to human interactions whereby saliva could be collected cautiously without immobilization. Additionally, Pearson was given skin nodule biopsies or saliva collected previously from six wild *L.africana* during GPS collaring operations by South Africa National Parks Veterinary Unit. All samples were shipped or hand-carried to USA with appropriate CITES permits.

### Search for evidence of EEHV and EGHV in Saliva from wild *L.cyclotis* in Gabon

While unable to personally collect samples from *L.cyclotis* in equatorial Africa, Pearson imported in 2017 saliva swab samples from 25 *L. cyclotis* that had been immobilized for GPS collaring purposes, courtesy Agence Nationale des Parcs Nationaux and CENEREST National Centre for Scientific and Technological Research, Gabon. The authors are unaware of any reports of skin nodules on wild *L. cyclotis*.

### Search for evidence of EEHV and EGHV in Saliva from seven wild-born *L. africana* at Six Flags Safari Park in USA

Concurrently, we investigated the frequency and types of EEHV and EGHV that were shed in saliva over an extended period of time (seven years) in saliva from the same individual *L. africanas.* In 2012, Pearson collected weekly saliva swabs from 2 adult female *L. africana* living in a herd of seven at Six Flags Safari Park, USA, and periodically collected saliva from the entire herd through 2019 for a total of 180 saliva samples. The two weekly collection candidates were the herd matriarch who had been imported from Uganda 1971-72 and had lived at the park ever since and a survivor of the ninety-nine *L.africana* imported from Zimbabwe in 1982-84 who had lived at several other USA zoos before coming to Six Flags in 2010.[60] The remaining five *L. africana*, imported with the matriarch from Uganda in 1971-1972, also had been at Six Flags continuously. Using many of the same primer sets, we detected many of the same species and subtypes of EEHV and EGHV that we found in wild *L.africana* and *L. cyclotis.* It is important to note that all Six Flags *L. africana* were asymptomatic for EHD at the onset and throughout this study. Our novel technique of collecting saliva by extra long swabs was equally effective to sample anaesthetized wild elephants or elephants such as these who were not accustomed to routine trunk wash or blood collection protocols used in many USA zoos to detect systemic EHD viremia. [63–68]. One caveat of this seven-year saliva study is that we have no saliva samples from any of the herd members prior to the introduction of the Zimbabwe-born *L africana* to address the question of viral cross-transmission within the herd.

Although not as readily subject to quantitative analysis (because of considerably more variability in individual elephant and handler collection techniques and recovered saliva volumes), in a similar study, this saliva swab plus Sanger PCR subtype DNA sequencing approach proved to closely parallel the results obtained in TW procedure for detecting EEHV3B in a young USA zoo *L. africana* recuperating from EHD [35]. The relative abundance of viral DNA obtained by the two approaches was not addressed but scoring of positive results seemed to be at least equally sensitive with both methods being able to detect the continued presence of viral DNA out to about eight months. The results also demonstrated that no significant change occurred in the EEHV3B DNA sequence obtained at each of three PCR loci over this post-viremia monitoring period. Further investigation of saliva using quantitative polymerase chain reaction (qPCR) is needed to evaluate saliva as a useful diagnostic and prognostic tool for early indication of EHD such as are blood and trunk wash. [69,70]. (**Figure 10**)

**Figure 10.**
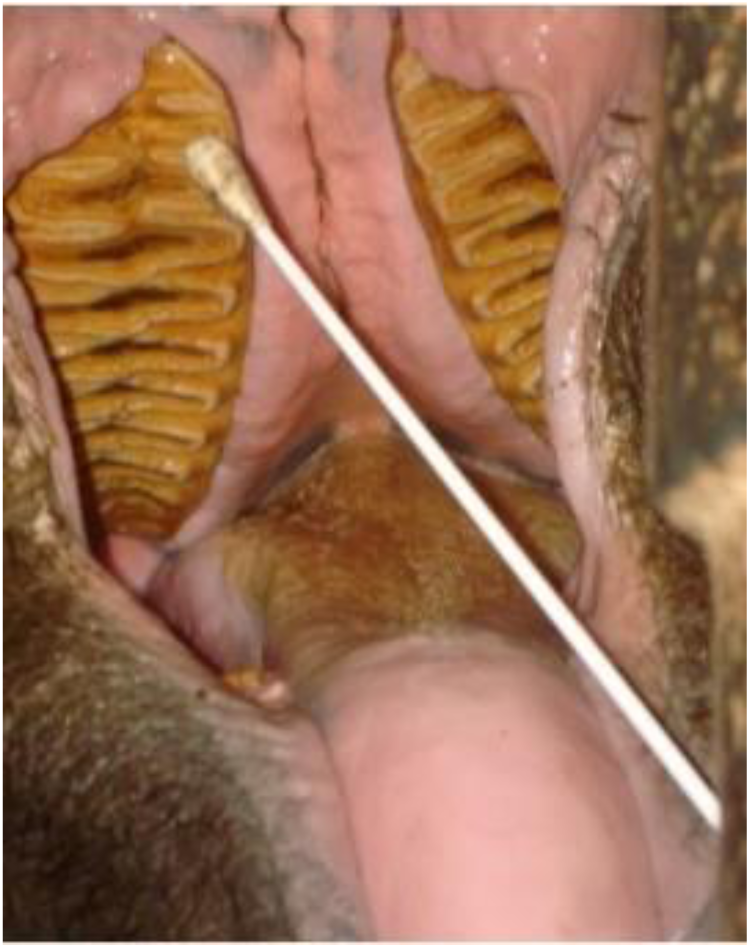
*L.africana* SFJ, Six Flags Safari Park, USA, 2012. Puritan 16” saliva swabs were collected by Holloway at Six Flags and Pearson. Credit Pearson, 2012.

## GENETIC EVALUATION

Pearson worked with these samples first at Enquist Laboratory, Princeton University, Princeton, NJ, USA, and subsequently at Rall Laboratory, Fox Chase Cancer Center, Philadelphia PA USA. Neither lab had previously ever worked with elephant herpesviruses. Because DNA contamination is very common in laboratories working with multiple biological samples at the same time, we exercised great caution in working with these samples, always using negative controls in every conventional polymerase chair reaction (cPCR), periodically testing our reagents for DNA contamination, and rerunning any nested cPCR that showed false positives in the negative gel electrophoresis lanes in separate laboratory space with fresh reagents. DNA was amplified by cPCR using three-round nested primer sets targeting many *Loxodonta*-associated EEHV genes loci including U38(POL), U39(gB), U47(gO), U66(TERex3), U77(HEL), U48.5(TK), U51(vGPCR1, U71/gM, U73(OBP), U77(HEL), U81(UDG), U82(gL), E24(vOX2B1), E4(vGCNT1), E37(ORF-0ex3), E9A(vOGT), E16D(vECTL), E54(vOX2-1), U14, U42(MTA) and U43(PRI) and primers designed specifically to recognize *Loxodonta-*associated EGHV genes loci. Many other primer combinations were used to distinguish unique polymorphisms of subtypes EEHV3A-EEHV3H, including chimeric domaines CD-I U39(gB), CD-II U47(gO), U48.5(TK), U48(gH), CD-III U81(UDG), U82(gL) E37(ORF-Oex3) CD IV E9A(vOCT) CD-V E16D(vECTL), among others. Sanger sequencing of purified DNA from nearly all the samples yielded unambiguous positive genetic matches to previously known *Loxodonta*-specific EEHV2, EEHV3A,EEHV3B EEHV6, EEHV7A, and EGHV1B, EGHV2, EGHV3B, EGHV4B, EGHV5B and discovered new rare species EEHV3C-H and EEHV7B and a novel subtype EGHV5A in the spleen of a zoo *E. maximus.* Many of the primer sets used could also have detected known *Elephas*-specific EEHV1A, EEHV1B, EEHV4, and EEHV5 if present, but they did not. For independent DNA extraction and cPCR and Sanger sequence verification, a subset of skin nodule biopsies sent to the Hayward Laboratory, Johns Hopkins University School of Medicine, yielded identical results. **Figure 11**, **Figure 12**.

**Figure 11.**
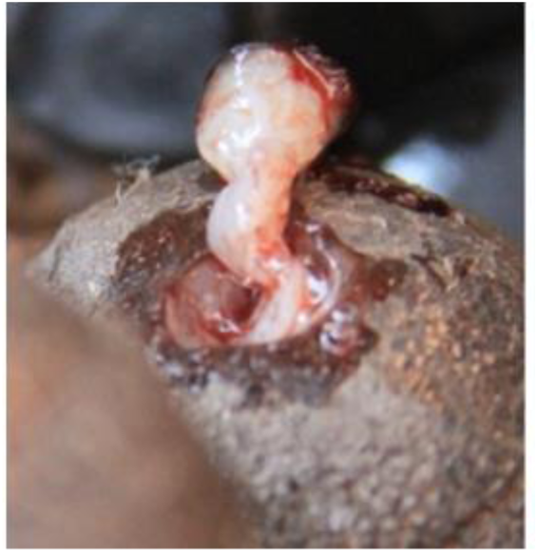
6 mm punch biopsy of skin nodule BW1M NOD1 positive for EEHV3A, EEHV3B, EEHV3D, EEHV7A and EGHV1B, Chobe National Park, Botswana, was immersed in Qiagen PAXgene Tissue Container, Credit Pearson 2013.

**Figure 12.**
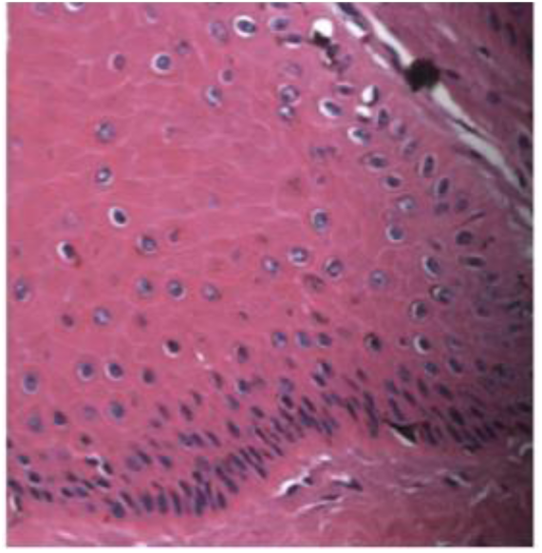
section of same biopsy BW1M NOD1 was fixed in formalin-fixed paraffin preserved (FFPE) block, cut at 5 microns and stained with hematoxylin and eosin. Histological tissue slide shows multiple keratocyte nuclei that are expanded by a dark basophilic material (intranuclear inclusions indicative of viral infection) distorting the nuclei into an oval to rounded appearance surrounded by a 1-2 um clear halo, mag 45x Credit SY Long, Hayward Laboratory.

**Figure 13.**
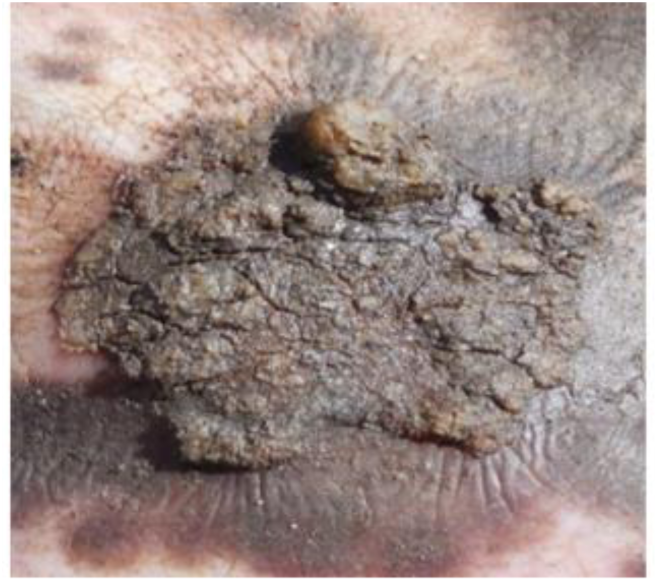
5 mm punch biopsies of ear wart RSA Wild Spirit positive for EEHV3A and EGHV3B, Balule, South Africa, was preserved in Qiagen RNAlater. Credit Pearson, 2013.

## COROLLARY STUDIES

In our efforts to understand the pathology of EHD, Pearson established elephant primary cells lines from elephant umbilical cords and amniotic sacs for culture and investigation of EEHV *in vitro.* One such cell line derived from the umbilical cord of a young *L. africana* born in a USA zoo that subsequently died of EEHV was used by the Vertebrate Genomes Project for assembly of the high-quality reference genome for *L. africana* [71], and others of these cell lines have contributed to the development of potential therapeutics to combat excessive inflammatory response to EEHV in *Loxodonta* and *Elephas* with Huntsman Cancer Center, Salt Lake City, UT USA [72]; and to the development of elephant pluripotent stems cells with Colossal Biosciences, Dallas, TX USA [73]. As previously published, Pearson et al 2021 used an isolate of African Elephant Polyomavirus (AelPyV-1) extracted from the same skin nodule biopsy from a *L. Africana,* BW1M Nod1,in Botswana, (Figure 10), that had tested positive for EEHV3A, EEHV3B, EEHV3D, EEH7A and EGHV1B, EEHV2 to create a recombinant plasmid, AelPyV-1-LTag, that can facilitate durable transformation of these primary elephant endothelial and epithelial cell lines.[74]. The finding in this corollary study that AelPyV-1 evidently is restricted almost exclusively to skin nodules suggest that further investigation is needed to ascertain whether AelPyV-1 or infiltrating lymphocytes latently infected with EEHV and/or EGHV may be responsible for the skin nodules such as we found in wild *L. africana*.

## METHODS AND MATERIALS

### Sample collection

Field sample collections were conducted under authorization of Elephants Without Borders Botswana Research Permit/ #WT8/36/4XV(41 (GPS-coordinates Chobe National Park, Kasane, Botswana, 17 °47’42.83”S, 25 °10’16.067”E); Kenya Research Permit #NCST/RRI/12/1MAS/9 Permittee Virginia R. Pearson; Save The Elephants Research Permit with Kenya Service Veterinary Capture and Services Department (GPS-coordinates Samburu National Reserve, Samburu County, Kenya, 0 °37’5”N, 37 °31’48”E and Keekorok, Masai Mara National Reserve, Narok, Kenya, 1 ° 35’ 9.00”S, +35 ° 15’ 6.00”E); and Elephants Alive with South African National Parks Veterinary Wildlife Services, (GPS-coordinates Timbavati Reserve, Mpumalanga Province, South Africa, 24 °20’07S,31 °20’38”E, and Crocodile Bridge, Kruger National Park, South Africa, 25 ° 21’0”S,31 °53’32”E); and CENEREST(National Centre for Scientific and Technological Research) and Agence Nationale des Parcs Nationaux Gabon, (GPS-coordinates Minkebe NP 1 °40’47”N,12 °45’34”E; Mwagna NP 0 °36’N,12 °42’E; Ivindo NP 0.088 N°, 12.63 E°; Pongara NP 0 °07’N,9 °38’E; Loango NP 2 °10’S,9 °34’E; Moukalaba Doudou NP 2 °26’S, 10 °25’E). Samples were imported into USA under authorization of United States Veterinary Permit for Importation and Transportation of Controlled Materials and Organisms and Vectors #11798, Permittee Virginia R. Pearson; San Diego Zoo Global CITES/ USFWS Import Permit # 13-US727416/9 and #10-US727416/9; International Elephant Foundation CITES/USFWS Import Permit #17US09806C/9; Botswana CITES Export Permit #0202118; Gabon CITES Export Permit #0785; Kenya CITES Export Permit # 008624/ OR82872; and South Africa CITES Export Permit #1087577. Protocols for elephant sample collection and laboratory analyses were approved by San Diego Zoo Global Institutional Animal Care and Use Committee IACUC #11–002; Princeton University IACUC Enquist Laboratory Protocol #1819; and Fox Chase Cancer Center Rall Laboratory federal regulations governing research involving material of animal origin (elephant). Samples from within the United States were collected according to participating zoos’ Animal Welfare and Research Committees and Association of Zoos and Aquariums Elephant Research and Necropsy Protocol.

### Immobilization and Biopsy collection

Elephants were immobilized by qualified, experienced field veterinarians using etorphine hydrochloride (M99). 5 mm punch biopsies were excised from trunk nodules and immersed in 2.5 ml RNALater Reagent (Qiagen). Incisions were treated with iodine and oxytetracycline. Diprenorphrine hydrochloride or naltrexone reversed sedation within 90 sec. Saliva samples from immobilized, or semi-captive wild or captive elephants in Africa and America were collected using 16 in polyester swabs (Puritan Medical Corporation) until saturated then immersed in 2 ml RNAprotect Cell Reagent or RNALater (Qiagen) or pressed onto Gentegra DNA Collection GenSaver Cards.

### DNA Extraction

DNA was extracted from skin nodule biopsies, saliva, and tissues using Qiagen DNeasy Blood and Tissue Kit according to the manufacturer’s instructions with the modification of extended overnight incubation at 56’C, followed by 10 min incubation at 95 °C. Archived (FFPE) biopsies, were re-embedded in fresh paraffin after which 8 mm microtome slices per biopsy were incubated in xylene at room temperature for 5 min shaking and rinsed with 100% EtOH a total of five times, prior to DNA extraction. Qiagen Repli-G was used to amplify extracted DNA in cases when three-round nested cPCR was necessary to detect expected very low levels of virus.

### Conventional Polymerase Chain Reaction (cPCR) see SI for primer list

For each 25 µl cPCR reaction, we used 2 µl elephant DNA, 12.5 µl GoTaq G2 Hot Start Green Master Mix (Promega), 8 µl sterile nuclease-free H2O, and 1.25 µl each forward and reverse primers. Reactions were run on an Eppendorf MasterCycler or Finnzymes PIKO Thermocycler programmed at 95 °C for 2 min; 45 cycles of 95 °C for 40 sec, 50 °C for 45 sec; 73 °C for 60 sec; final extension 78 °C for 7 min. 25 µl of each PCR reaction was run on 2% agarose electrophoresis gels with ethidium bromide or GelRed or GelGreen nucleic acid stain (Phenix Research Products). Bands of expected size were purified and sent for sequencing.

### Sequencing

Primer extension sequencing was performed by Genewiz, Inc (South Plainfield, NJ) using Applied Biosystems BigDye version 3.1. The reactions were then run on Applied Biosystem 3730xl DNA Analyzer.

## RESULTS

### Evidence of EEHV and EGHV in skin nodules and saliva from wild *L.africana* in Kenya, Botswana, South Africa and Zimbabwe,

DNA sequences of multiple species of EEHV and EGHV were identified in skin nodule biopsies from one young female calf (Samburu NP, Kenya) and one male calf (Maasai Mara NP, Kenya) with two nodules each; two young female calves (Samburu NP, Kenya and Chobe NP, Botswana, with one nodule each; and one young calf male with 4 trunk nodules (Chobe NP, Botswana); and 4 sub-adult females (2 Maasai Mara NP,. Kenya, and 2 Kruger NP South Africa) with one nodule each; in ear wart-like lesions from an adult South African bull *L. Africana*, (Timbavati, South Africa); and in random necropsy lung tissue from a sub-adult female *L. africana* in Samburu, Kenya, which Pearson et al previously reported[44–48]. These were selectively screened using combinations of up to 21 different cPCR primers (including first, second and third round amplification for each when appropriate) that detected known selected high GC-rich *EEHV3* or *EEHV7* loci, as well as several more generic cPCR primer sets that also detected *EEHV2* or *EEHV6*. 100% of nine (n=9 skin nodule biopsy and one (n=1) ear wart biopsy collected from wild *<u>L.. africana</u>* adults and juveniles from Kenya (5), Botswana (2) and South Africa (3) were found to be low level positive by nested cPCR and Sanger sequencing for EEHV2, EEHV3A, EEHV3B, EEHV6, EEHV7A and EEHV7B and a wide variety of subtypes or strains, including novel subtypes of EEHV3B and EEHV7B and rare subtypes EEHV3C to EEHV3F as well as multiple examples of EGHV1B, EGHV2, EGHV3B, EGHV4,EGHV5BEGHV3B, EGHV4B and EGHV5B. (**Table 2**.)

**Table 2.**
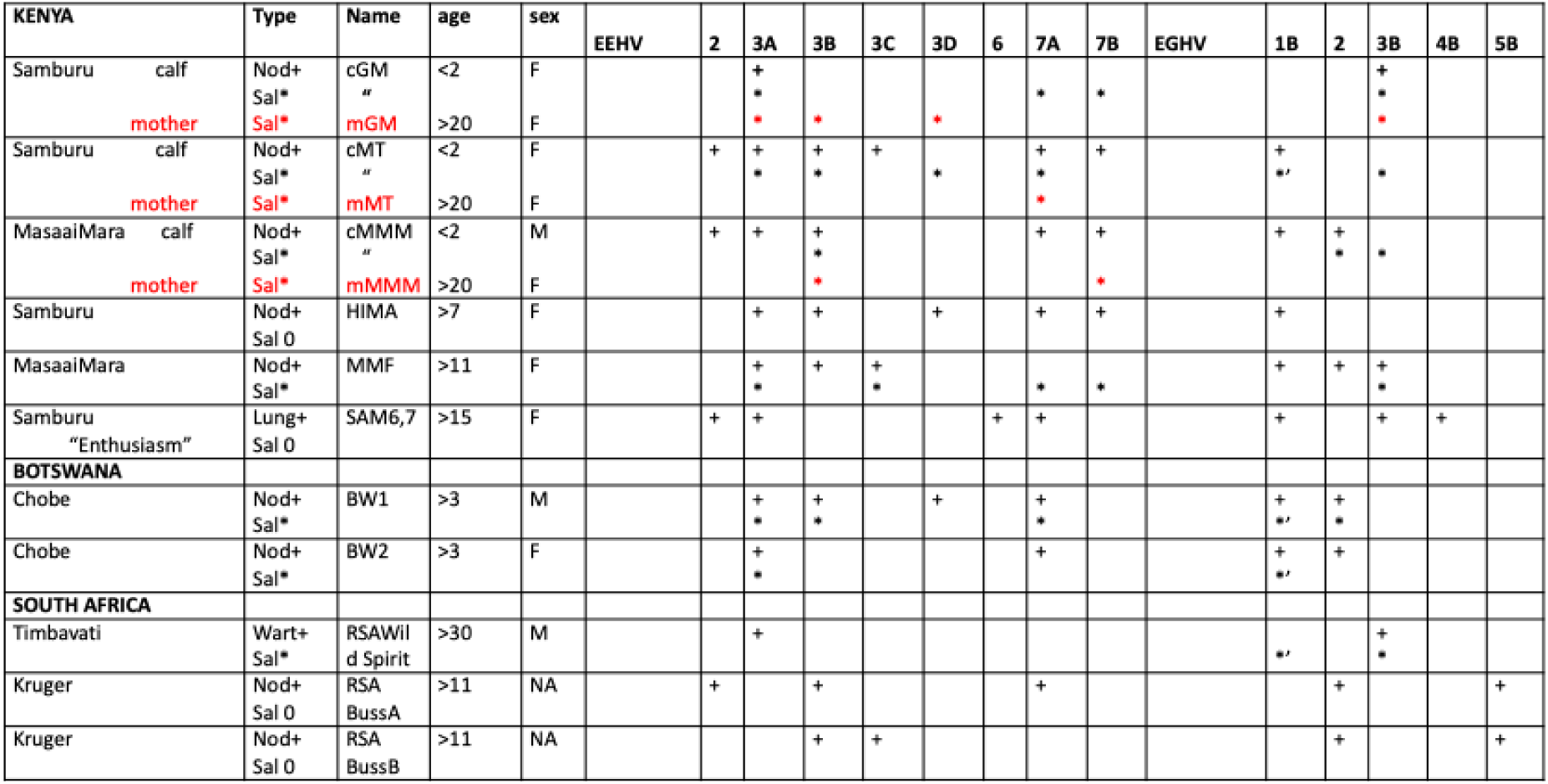
EEHV and EGHV in skin nodules (including matching saliva samples) and warts.

Saliva from Kenya and Botswana mother plus calf pairs(n=6, positive 6/6=100%) and from 34 (n=34, positive 29/40=73%) additional *L.africana* in Botswana, Kenya and South Africa and Zimbabwe were screened with up to seven selected GC-rich branch cPCR loci (POL, HEL, U71/gM, OBP, TK, vGCNT1 and UDG) yielding more than 75 positive unambiguous EEHV DNA sequences of EEHV2, EEHV3A, EEHV3B, EEHV3C, EEHV 3D, EEHV6, EEHV7A and EEHV7B and an additional two dozen unresolvable mixtures. Saliva samples from four of the same Kenya juveniles with nodules also detected between one and four subtypes of EEHV3A, EEHV3B, EEHV3C, EEHV3D and one or two subtypes of EEHV7A and EEHV7B as well as EGHV1B and/or EGHV3B each, some of which matched the strains in the skin nodules but mostly did not. Saliva from some mother/calf pairs were identical, but more often different. (**Table 3**.)

**Table 3.**
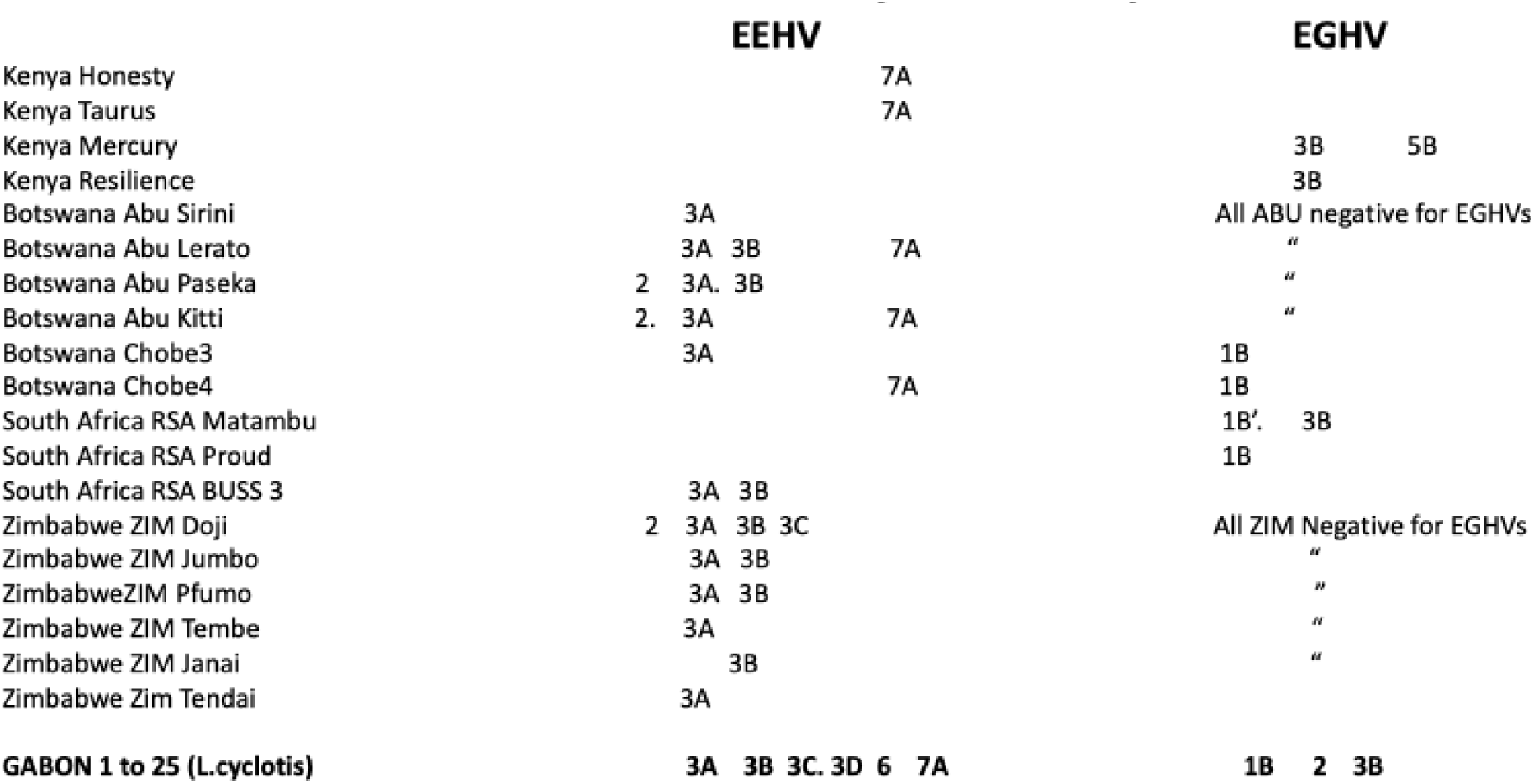
EEHV and EGHV in Saliva from *L. Africana and L.cylotis* in Africa.

### Evidence of EEHV and EGHV in Saliva from wild *L. cyclotis* in Gabon

Saliva swabs from 25 *L. cyclotis* were screened by cPCR and Sanger sequencing with more than six GC-rich branch EEHV primer sets including POL, HEL, TER, UDG, TK and OBP loci. Over half of 25 saliva samples from *L. cyclotis* in Gabon were positive for at least one *EEHV3* or *EEHV7* strain, and once for a novel *EEHV6*, but no *EEHV2* was detected, and many of these were also present as unusual, rare variants of non-A/non B Subtypes of *EEHVs* that we found in nodules and saliva from *L. africana.* In terms of number of distinct virus strains in individual samples, five were negative, 8x had just one strain, 8x had two strains, 3x had three strains, 1x had four strains and 1x had five different strains present. Unusually, unlike the results from the nodules, only a single EEHV strain was detected in any one saliva sample. The high prevalence of EEHV3B suggests that most likely EEHV3B and the more rare EEHV3C, EEHV3D, EEHV3E, EEHV3F subtypes have been exchanged and recombined during overlapping range interactions between *L.africana* and *L. cyclotis* hosts. These results closely resemble previous published findings by Zong et al 2015 of EEHV in lung nodules and necropsy lung tissue [43]. Overall, the predominant result was for an EEHV3A strain (21 positive/43 tested=49% of examples), whereas EEHV3B strains were detected 17 of 43=39.5%, EEHV2 strains three times (just with the POL primers) 7/43=16%and EEHV7A three times (HEL or U71/gM.) 7A=10/43=23%, 73/7%. We attribute the lower percentage of positives (n=25, positive 13/25=52%) to insufficient volume of saliva collected in the swabs. (**Table 4**.)

**Table 4.**
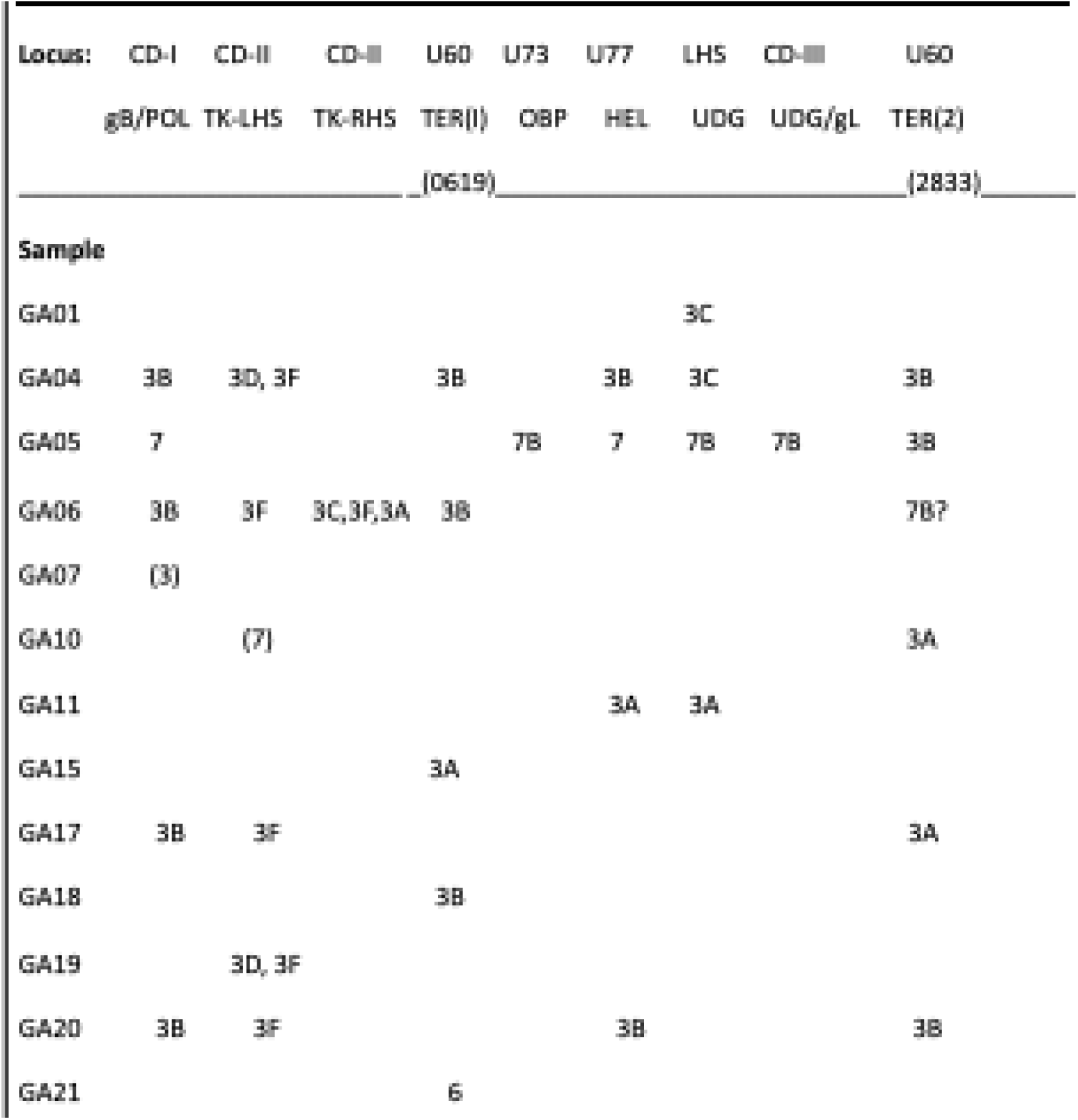
All Gabon Loxodonta cyclotis OCR Sequence Data (Subtypes at 8x Loci.

### Evidence of EEHV and EGHV in saliva collected over seven years from *L.africana* in seven wild-born *L.africana* at Six Flags Safari Park, USA

Our temporal saliva study of seven *L. africana* at Six Flags Safari Park, NJ, USA, showed that EEHV2, EEHV3A, EEHV3B, EEHV6, EEHV7A and multiple EGHVs are periodically shed in saliva, with frequently more than one virus being detectable at a time. Primers used were TER, POL, OBP and UDG. Saliva from two of seven adult females, SFL and SFJ, were collected approximately weekly over a full 12-month time period from January, 2012 through January, 2013 for a total of 43 samples each for Year One (Y1) and thereafter 12 times until through 2019 for a total of 55 samples each. The other five were samples twice during the 2012 and thereafter a total of twelve times for a total of 14 samples each, for an overall total of 180 saliva samples. Overall, SFJ scored positive for EEHV2 in 17/43=40% of Y1 samples while SFL scored positive for EEHV2 in 8/43=19% of Y1 samples In contrast, SFJ was positive for EEHV6 in 17/43=40% Y1 samples and SFL was positive for EEHV6 in 15/43=35% of Y1 samples. Both SFL and SFJ shed both EEHV3A and EEHV3B intermittently for a total of SFJ 11/ 43=26% and SFL 10/43=23%. SFJ was positive four times for four different viruses at the same time (EEHV2, EEHV3A, EEHV3B and EEHV6) whereas SFL was positive at some point for two or three different EEHV species at the same time (either EEHV2, EEHV3B and EEHV6 or EEHV2, EEHV3B).Overall, among seven adult *L. africana* tested at Six Flags Safari Park from 2012 to 2019, 6/7=86% were positive for EEHV2, 5/7=71% were positive for EEHV6, 4/7=57% for EEHV3B, 4/7=57% for EEHV3A and 2/7=29% were positive for EEHV7A. Only a single sample from SFJ, EEHV2 POL (SFJ #4, January 2012), gave a band of sufficient intensity to obtain clear unambiguous DNA sequence data directly from the first round cPCR amplification. All others required second or third round amplification first. We interpret this as evidence of previous persistent infections rather than necessarily currently active systemic infections or shedding as might be detected in the first round cPCR of EHD cases. (**Table 5**.)

**Table 5.** EEHV and EGHV in Saliva in L. africana at Six Flags Safari Park, NJ, USA.

| Name | Host | Loc | EGHV1 |  |  | Other EGHV |  |  |  |  |  | EEHV |  |  |  |  |  |
| --- | --- | --- | --- | --- | --- | --- | --- | --- | --- | --- | --- | --- | --- | --- | --- | --- | --- |
|  |  |  | TTTT | CCGC | 1A | 2 | 3A | 3B | 4 | 5A | 5B | 2 | 3A | 3B | 6 | 7A | 7B |
| SFJ | *L. africana* | USA | 1B |  |  | 2 |  |  |  |  |  | 2 | 3A | 3B | 6 |  |  |
| SFL | *L. africana* | USA | 1B |  |  |  |  |  |  |  |  | 2 | 3A | 3B | 6 |  |  |
| SFT | *L. africana* | USA |  | 1B' |  | 2 |  |  |  |  |  |  | 3A |  | 6 |  |  |
| SFG | *L. africana* | USA |  | 1B' |  |  |  |  |  |  |  | 2 |  |  | 6 | 7A |  |
| SFS | *L. africana* | USA |  | (1B') |  |  |  |  | 4 |  |  | 2 |  | 3B |  | 7Anovel |  |
| SFD | *L. africana* | USA |  | 1B' |  |  |  |  |  |  |  | 2 | 3A |  | 6 |  |  |
| SFB | *L. africana* | USA | 1B |  |  |  |  |  |  |  |  | 2 |  | 3B |  |  |  |

### Types of EGHV detected in *L. africana* and *L.cyclotis*

Sixteen examples of EGHV1B (the most abundant EGHVs species we encountered) were detected within sub-Saharan Africa, including two invariant polymorphism patterns that differ by only 4/450-bp (0.9%). The more commonly found TTTT pattern was seen in 13 of 14 EGHV1B cases from *L.africana* within Africa, while the more rare CCGC pattern was found only once in *L. africana* adult bull Matambu (SAG5) (Timbavati, South Africa) in continental Africa. All three EGHV1B positive samples from *L. cyclotis* fall into the same CCGC variant suggesting distinct host species origins for these two variants. These same two polymorphic variants were detected in three whole herd saliva sets from Six Flags Safari Park. USA SFJ, SFL, SFG, SFS and SFB carried the EGHV1B TTTT variant whereas SFT and SFD carried the CCGC variant identical to South African RSA Matambu (SAG5) as noted by asterisk* in TABLE 2. SFJ saliva sample #97 collected on November 21.2012 has been designated the prototype for EGHV1*B*, NAG9, Genbank #KC810011.1. In saliva samples from additional USA *L. africana* (not shown) we only found the CCGC variant in less than one-quarter of the samples tested.

EGHV2 was detected in POL locus in skin nodules from wild *L. africana* RSA BussA, KENYA MMM, KENYA MMF, and Botswana BWM1, BWF2 and in saliva from Six Flags Safari Park USA SFJ and USA SFT which are identical to those EGHV2 sequences reported in a 38 year old zoo *L. africana,* Genbank acc# KU726836, # KU726837 [35] and were just one to four bp different from the prototype *E. maximus* EGHV2 POL.

EGHV3B was detected frequently in skin nodules, ear warts and saliva from *L. africana* and in saliva from *L. cyclotis*. Quite unexpectedly however, we detected widespread presence of EGHV3B in necropsy organs from an adult female *L. africana,* NAG31, in a USA zoo, including in ear lesions, heart, lymph node salivary gland, serum and thymus. We have rarely detected any EGHVs in any internal organ tissues from *EHD* cases or necropsy samples in USA zoos. Surprisingly, no AelPyV-1 DNA was detected in the ear lesions. (**Figure 14**.)

**Figure 14.**
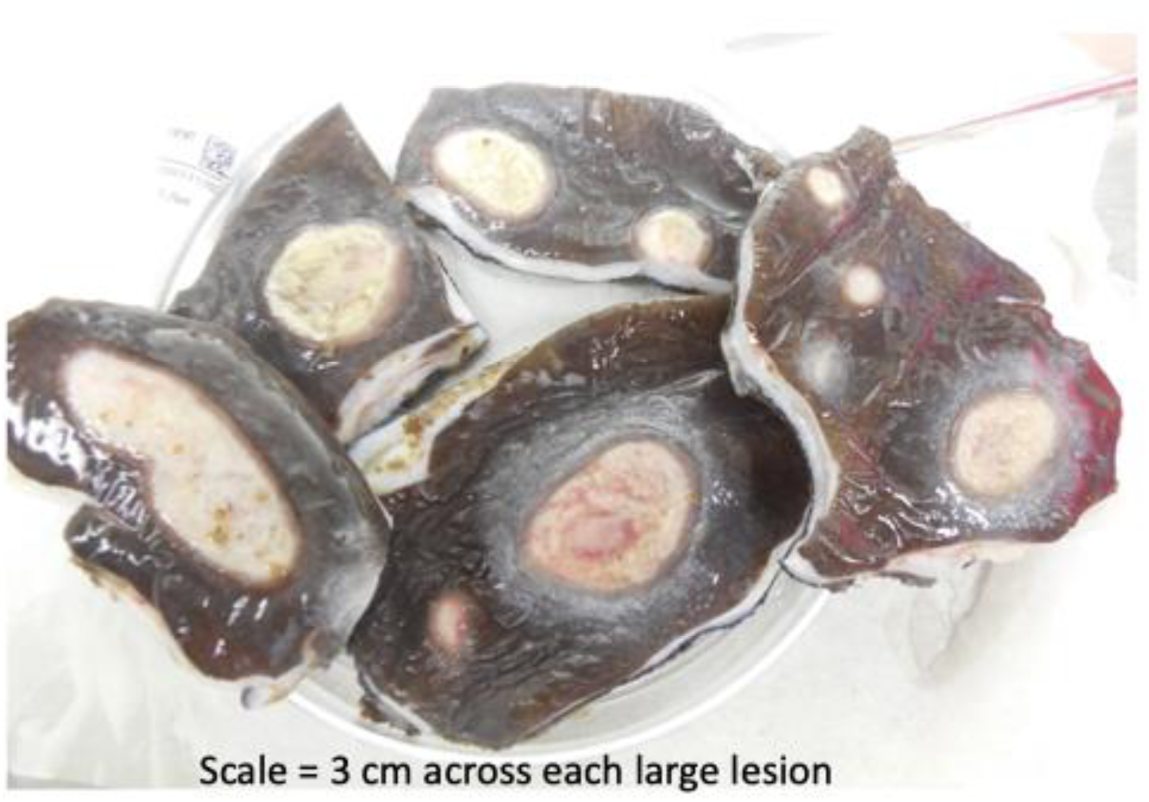
Ear lesions on back of ear from necropsy samples of adult female *L. africana* NAG31 in USA zoo, received by Pearson, 2014. Ear lesions, heart, lymph node salivary gland, serum and thymus, were positive for EGHV3B only, no EEHV was detected. Credit Pearson, 2014.

EGHV4 was detected in saliva from SFS at Six Flags Safari Park; in the young zoo *L.africana* that survived EEHV3B [25]; and in wild *L. africana* lung biopsies designated KENYA Sam 6 and Sam 7 that Pearson collected in 2011 [43–47]. Subsequently, we discovered an evidently novel Asian elephant version of EGHV4 in spleen tissue from a three year old male captive-born *E.maximus* NAG72 in New Mexico, USA, that died suddenly of EHD, Genbank Acc. #PQ379918. Identical 250 bp sequences were obtained from saliva from a captive-born living male juvenile *E. maximus* NAG71 in Texas and in the spleen from a deceased elderly USA *E. maximus* NAG56 in Florida, USA. These results led us to reassign all the previous *Loxodonta-*associated EGHV4s and *Elephas*-associated EGHV4*s* as distinct subspecies EGHV4A or EGHV4B endogenous to either *Elephas* or *Loxodonta* respectively.

EGHV5B was detected in skin nodule biopsies from two sub-adult *Loxodonta africana* in Kruger National Park, South Africa: BussA SAG3, BussB SAG4, and in saliva from an adult femal*e L. africana* in Kenya designated KENYA Mercury KG16. In this case, the 450-bp DNA sequences obtained after second round PCR with the generic Pan-EGHV primers were identical to one another and to those from the prototype EGHV5B (Genbank accession #JX268522 [55]. Most recently, Sanger sequencing from first round cPCR using *EGHV* PAN POL primers R1/L1 of a skin nodule biopsy from the trunk of a 3 year old captive-born *L.africana* in England yielded a 439 base pair sequence of EGHV5B, Genbank Acc# PQ3979917, that is identical to the prototype NAG8. The sequence obtained from the R1 and L1 primers, was round 1 PCR, R1 LGH6784B 5’-GTGGTKGACTTTGCYAGCCTSTACCC-3’ L1 LGH7489 5’-GTCRGTGTCYCCGTAGAYNAC-3’. This is the first time we have been able to compare EGHV sequence results from a skin nodule on a *L. africana* residing outside of Africa to skin nodule biopsies and saliva collected in Botswana and Kenya. (**Figure 15**.)

**Figure 15.**
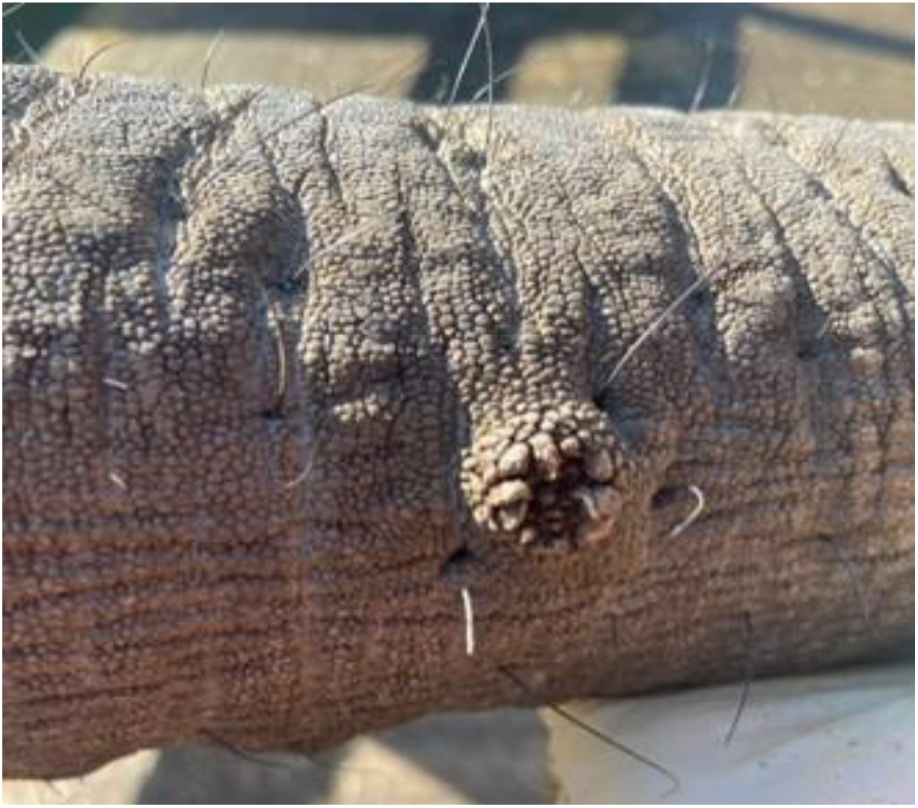
Trunk nodule on 3 year old *L. africana* positive for *EGHV5*,Credit J. Hopper, DVM, Howletts Wild Animal Trust; DNA extraction, cPCR and Sanger sequencing provided by A. Dasjerti, DVM, APHA, UK, for Howletts Wild Animal Trust, England, 2024.

## DISCUSSION

Nearly all skin nodules examined proved to be much more complicated, contained multiple different virus types and strains and gave at least one positive result with one or more of the primer pairs used. Furthermore, a large fraction evidently harbored multiple EEHV genome types and indeed unlike with any Asian elephant viremia, TW or saliva swab samples evaluated, many of the skin nodules gave mixed DNA sequences from multiple different genomes being present within the same purified PCR bands. This phenomenon of multiple simultaneously infecting genomes that sometimes gave mixed DNA sequence profiles was also seen in the earlier study with the much smaller number of African elephant lung nodules (Zong, 2015), but it has been exclusive to just the skin and lung nodules or to asymptomatic necropsy samples, but it has been very rare or non-existent within the saliva studies even from the same group of elephants. Note that some mixtures gave clean enough sequences that they could be resolved with or without additional nested PCR analysis, although many could not of course and the latter have just been recorded here as “mixed 3/7” or “mixed 3A/3B” or “multiple 3A” strains present, without accompanying Genbank DNA sequence files. However, wherever possible clean DNA sequence files if of sufficient length to reveal phylogenetically useful characteristic polymorphisms have been generated, edited and submitted to Genbank. A further rather curious feature of the results from the Kenya skin nodules was that no two nodules from the same animal seemed to have the same complement of virus species, subspecies or strains present and that the strains even of say just EEHV3A present in different nodules from the same animal often proved to be distinctly different with multiple novel polymorphisms present. Indeed, for EEHV3 especially, the large number of nucleotide variations found between the numerous different strains encountered and patterns of subgrouping among them have often allowed us to designate more than just EEHV3A and EEHV3B and further define subtypes EEHV3C, EEHV3D and EEHV3E that may contain up to ten localized nonadjacent chimeric domains (CD) or hypervariable gene blocks encompassing anywhere up to seven distinct subtype clusters with large (15 to 50%) protein level differences. EEHV3 strains fall into two major chimeric subtype clusters similar to those described previously in *Elephas* as EEHV1AB, EEHV4A/B and EEHV5A/B [53] In EEHV3A strains only, parts of the CDs including vECTL, gB. gH, TK, vGPCR1, gL and UDG can instead have several other alternative highly diverged but unlinked subtype clusters (designated EEHV3C, EEHV3D, EEHV3E, and EEHV3F, Genbank acc.# MN373268, NC077039, NAP42 EEHV7A Genbank acc# KU321582, NAP62 Samson EEHV3B KT832467.1-KT832476.1, KT832491.1-KT832495.1 []. These differ greatly (by between 15 and 45%) in several linked but nonadjacent chimeric domains (CD-I, CD-II, CD-III, and CD-V) totaling 15 to 20-kb in size, but by just 2 - 3% everywhere else. Our ongoing genetic evaluation of the few recent EHD cases attributable to EEHV2 and EEHV6 suggest that similar division into A and B subtypes may be appropriate, as found in EEHV3A/3B and EEHV7A/7B pairs that differ by 7 - 9% overall with many other genes not just the CDs varying significantly. (Genbank acc# NC077039, MN373268) [51,52]. Based largely on the summed higher levels of individual protein divergence across the whole EEHV3A versus EEHV3B genome as well as the relatively high frequency of EEHV3B being found in saliva swabs from Gabon, compared to EEHV3A being far more prevalent in Botswana, Kenya and South Africa we judge that these two EEHV3 groups should in fact be designated as distinct species with EEHV3A likely having evolved in *L. africana* and with EEHV3B probably having evolved originally in *L. cyclotis*.

In addition to the multiple EEHV subtypes associated with HD, most healthy asymptomatic adult elephants evidently harbor and periodically shed as well DNA from the second herpesvirus sub-family EGHV. Overall nodule biopsies from six of the seven wild *L. africana,* the KENYA SAM6/SAM7 lung biopsies, RSA Wild Spirit (SAG7) non-nodular ear wart biopsy and more than half of the saliva samples from 29 wild *L. africana* and 25 *L. cyclotis* and 70 *L. africana* from North America (additional NA samples not included in Table 3) proved positive for clean unambiguous detection of at least one type of EGHV, although many more clearly contained unresolvable mixture of multiple EGHV genome types. These studies have yielded unambiguous positive results for EGHV infections in more than 100 elephants out of 125 tested. In our *Loxodonta* saliva swab analyses, very few samples were positive directly on first-round cPCR, and most were only detected after either second-round or even third-round nested cPCR. these represent values of only a few 100 to a few thousand viral genome equivalents (vge) per ul/ug of DNA. In contrast, in the blood or tissue from severe HD cases the cPCR is usually still first-round positive even after 10,000-fold dilution of the DNA sample (representing many millions or even tens of millions of viral genomes (vge) per ml of blood or per ug of total DNA present. We interpret this as evidence of previous persistent infections rather than necessarily currently active systemic infections or shedding as might be detected in the first round cPCR of EHD cases. Therefore, these and similar results should never be interpreted as diagnostic of disease in an asymptomatic healthy elephant, but merely as evidence of a very low-level sporadic persistence following a previous infection that might or might not qualify as periodic reactivation from latency.

## CONCLUSION

Many of the primer sets used to detect multiple types of *Loxodonta-*associated EEHV in skin nodules, saliva and tissues from wild and zoo *L. africana* and wild *L. cyclotis* could also have detected *Elephas-*associated EEHV if present in our samples, but they did not. The unambiguous distinct types of EEHV we have identified in *Loxodonta* dispel the originally held hypothesis that the severe pathogenesis and mortality attributable to EEHV in *Elephas* might be caused by lethal cross-species infections from *Loxodonta*. We conclude that persistent low-level infections of EEHV2, EEHV3A, EEHV3B, rare subtypes EEHV3C-EEHV3H, EEHV6, EEHV7A and EEHV7B, and EGHV1B, EGHV2, EGHV3B, EGHV4B and EGHV5B are endemic viruses of *L. africana* and *L. cyclotis.* Our extensive library of EEHV and EGHV sequences from wild and zoo *Loxodonta* that has been archived in digital format at the International Elephant Foundation will be a significant contribution to the elephant virology community.

## ACKNOWLEDGEMENTS

We thank Lynn W. Enquist, PhD, Princeton University, USA; Glenn F. Rall, PhD, Fox Chase Cancer Center, USA; Jason Holloway, William Rives, DVM, Kenneth Keiffer, DVM, Six Flags Safari Park, USA; Michael Chase, PhD, Kelly Landen, Larry Patterson, DVM, Elephants Without Borders and Government of Botswana; Iain Douglas-Hamilton, PhD, David Daballen, Jerenimo Lepirei, Gilbert Sabinga, Chris Leadismo, Lucy King, PhD, Save The Elephants, and Mathew Mutinda, DVM, Domnic Mjele, DVM, Kenya Wildlife Service Veterinary and Capture Services Department, and Government of Kenya; Michelle Henley, Phd, Elephants Alive, South Africa; Peter Buss, DVM, South African National Parks Veterinary Service and Government of South Africa; Pete Morkel, DVM, Stephanie Bourgeois, PhD, Chimene Nze Nkogue, PhD, Agence National des Parcs Nationaux, CENAREST, and Government of Gabon; Elliott Jacobson, DVM, University of Florida School of Veterinary Medicine, USA; Oliver Ryder, PhD, Christina Reif, San Diego Zoo Wildlife Alliance, USA; Deborah Olson, International Elephant Foundation, USA; Jane Hopper, DVM, Howletts/Port Lympne, England; SY Long, Sarah Heaggens, Johns Hopkins School of Medicine, USA; Albuquerque Zoo, Dickerson Park Zoo, Caldwell Zoo, Cheyenne Mountain Zoo, Cleveland Zoo, Dallas Zoo, Denver Zoo, Fort Worth Zoo, Have Trunk Will Travel, Houston Zoo, Kansas City Zoo, Louisville Zoo, Oklahoma City Zoo, Oregon Zoo, Maryland Zoo in Baltimore; Nashville Zoo, Philadelphia Zoo, Pittsburgh Zoo, St. Louis Zoo and many others with elephants collections, USA; Heidi H.Robbins, Susannah Rouse, the Pearson family, Gentegra LLC, Qiagen Inc., Promega Corporation, Puritan Medical Products.

## Supplemental Information

**S1 PRIMERS**

## AUTHOR CONTRIBUTIONS

Sample collection, DNA Extraction, cPCR, gel electrophoresis, DNA purification: Pearson

Primer Design: Hayward

Sequencing Analysis: Hayward and Pearson

Original manuscript drafts: Pearson and Hayward

Review & editing: Pearson

