## Supplemental S1 for "Persistent Low-Level Infections of Elephant Endotheliotropic Herpesvirus and Elephant Gammaherpesvirus Detected in Skin Nodules and Saliva from Wild and Zoo African Elephants"

**Primers used in this study:**

**SET 1 U38 PAN POL**

R1 LGH6711 5’-GTATTTGATTTYGCNAGYYTGTAYCC-3’

R2 LGH6712 5’-TGYAAYGCCGTNTAYGGATTYACCGG-3	L1 LGH6710 5’-ACAAACACGCTGTCRGTRTCYCCRTA-3’	rd1 R1/L1 = 530-bp; rd2 R2/L1 = 250-bp.

**SET 2A U60 PAN TER**

R1 LGH6671B 5’-GTTYGTAGTAAANGCCGGATCTAC-3 ‘

R2 LGH3025B 5’-GGTACTATATCTTATCATRTC-3’

R3 LGH-R3alt	5’-GGGGTTTGCCAGGAAGCTAGTG-3’

L2 LGH-L2-2A 5’-GTAATCTCSTATGTGTGYGAGGAG,C-3

L1 5’ LGH7576B 5’-GCNCACGTRAAGAARACACCG,AC-3’ rd1 R1/L1 = 360-bp, rd2 R2/L1 = 290-bp, rd3 R3/L1 = 210-bp.

**SET 4 U71/(gM)**

R1 LGH6749	5’-CTATGGGATCCGAACTTTC-3’

R2 LGH6750	5’-CTTTCTAAGGGGGTTTGTTGC-3’

R3 LGH6793	5’-GCCATCGGGATCCGGAAAACGCC-3’

L3	LGH6792	5’-GGGCATCTTCTTCTCCCTCATAGC-3’ L2	LGH6752	5’-CTACATGCCCATGCAGATAGG-3’

rd1 R1/L1 = 750-bp, rd2A R2/L1 and rd2B R1/L2 =730-bp, rd3 R2/L2 = 710-bp,

alt rd 2B R1/L3 = 680-bp and alt rd2C R3/L1 = 700-bp; **SET 5 U51 (vGPCR1) specific for *EEHV2* (not good for *EEHV5*)**

R1 LGH7506	5’-GATTGTGAACGCTGTATGTC-3’

R2 LGH7470B	5’-GACAGGTGGTACTGTATGATGTGC-3’

R3 LGH7471	5’-CGGTTACACCGTACCGTGGCTTGC-3’

L3 LGH5201	5’-GCCAGGGTAGATAGAATCAAGGGAA-3’

L2 LGH5200A	5’-CGTGATACGCTTCCAAACATACAGC-3’	L1 LGH4963B	5’-GACTTTCTTCGTGTAGCCCTCGTCTT-3’

rd1 R1/L1 = 910-bp, rd2A R1/L2 = 750-bp, rd3A R1/L3 = 690-bp, and rd2B R2/L1 = 730bp, rd3B R3/L1 = 550-bp or R2/L2 = 570-bp.

**SET 5B U51(vGPCR1) specific for *EEHV1* only (outside pair OK for *EEHV6*)**

R1 LGH9323	5’-GTGCTAAACCTTTTCAACGAGACTTC-3’

R2 LGH9351	5’-GCGTTTAACGCMACRCAGTTGGT-3’

L2	LGH9353	5’-CGTAACAGGTTAGCATGAGTATCCTCTTCC-3’	L1	LGH9325	5’-GAGAACGCGTTCTGACTTTCTTCATCA-3’

rd1R1/L1 = 1100-bp; rd2 2A PCR R1/L2 = 990-bp, rd 2B PCR R2/L1 = 1070-bp; rd3AB R2/L2 = 960-bp.

**SET 6 U77 PAN HEL**

R1 LGH6649 5’-CCAGTCAACGTATAGCTCGTAG-3

R2 LGH6743 5’-GCAAGGTRGAACGTATCGTCG-3

L3 LGH7990 5’-CACCCACCCARTTGTAGGGAAAGTGC-3’

L2 LGH7885 5’-CTGCGTGTAACATGTGTTC-3’

L1 LGH3198 5’-CACAGMGCGTTGTAGAACC-3’

rd1A R1/L1 = 980-bp, rd2A R1/L2 = 950-bp, rd2B R2/L1 = 685-bp, rd3A R1/L3 = 500-bp, rd3B R2/L2 = 540-bp

**SET 6B U77(HEL) specific for GC-rich *EEHV3/4/7***

R1-7’	LGH7920B 5’-CATGTTSAGGTAGTGCAC**C**GA**G**AGC-3’

R2-7	LGH7921	5’-CTGAACATGTGTTCGGCCAGCAAGG-3’

L2-7	LGH7922	5’-CAGCAGCTAGGAGACGTGGCGAC-3’	L1	LGH3198	5’-CACAGMGCGTTGTAGAACC-3’	rd1 R1-7’/L1 .900-bp, rd2A R1-7’/L2-7 = 500-bp, rd2B R2-7/L1 = 650-bp, rd3AB R2-7/L2-7 = 370-bp.

**SET 7A U60 TER specific for *EEHV3/4***

R1 LGH6707 5’-GTGCTGTAGCGGATCATGTC-3’

R2 LGH6727 5’-GCAACACGAGCACGCAAAGTACGTC-3’

L2 LGH6728 5’-CGGATCATGTCGAACTCCGTG-3’

L0 LGH-LO-7 5’-CGTCGAACACGAGCACGCAAAGTACGTC-3’

rd1 1A R1/L1 = 310-bp, rd2A 2A R2/L0-7 = 300-bp, rd2B R1/L2 = 300-bp, rd3A/B R2/L0-7 = 280-bp.

**SET 8 U38 POL specific for *EEHV2***

R1	LGH7440	5’-GACTTCGCCAGCTTGTATCC-3’

R2	LGH7437	5’-GTATCATCAAGCTTATAACC-3’

L2	LGH7450	5’-CTCTACATTTACCGTACACTC-3’ L1	LGH6525	5’-CACATCGATACGGAATCTC-3’	rd1 R1/L1 = 510-bp, rd2A R2/L1 = 490-bp, rd2B R1/L2 = 490-bp, rd3AB R2/L2=470bp.

**SET 8A U38 POL specific for *EEHV2/5***

R1	LGH8423	5’-CATATACGAATGTGCCTCRGAATAYGAGC-3’

R2	LGH8424	5’-GACATGTATCGYGTGTGYATGGATAAGG-3’

L2	LGH8425	5’-CCCGTGTTAGTGGTGACAGTCGTAAC-3’	L1	LGH8426	5’-CCGCCAACCAGGATGTAAGAAGTTG-3’

rd1 R1/L1 = 890-bp, rd2A R1/L2 = 630-bp, rd2B R2/L1 = 555-bp, rd3AB R2/L2 =450-bp.

**SET 9 U38 POL specific for *EEHV3/4***

R1	LGH7400	5’-CAGCATCATCCAGGCCTACAAC-3’

R2	LGH6720	5’-ATCCTGGCGCAGCTGCTGAC-3’

R3	LGH6721	5’-CTCACCTGCAACGCCGTCTA-3’

L1	LGH6719	5’-CGTTGAAGGTGTCGCAGAT-3’

R1’ LGH8750	5’-CTGCTACTACTCCACCCTCGTSCTGG-

L1’	LGH8751	5’-GCCGTGACGGACTCGGCCACGGCCAG-3’

rd1 R1/L1 = 390-bp, rd2 R2/L1 = 270-bp, rd3 R3/L1 = 15-bp, and alt rd2A R1’/L1 = 360-bp, alt rd2B = R1/L1’ = 330-bp, alt rd3A R1’/L1’ = 300-bp

**SET 12A *EGHV* PAN POL (first step)**

R1	LGH6784B 5’-GTGGTKGACTTTGCYAGCCTSTACCC-

R2	LGH6785	5’-CCCMAGYATTATWCAGGCMCA-3’	L1	LGH7489	5’-GTCRGTGTCYCCGTAGAYNAC-3’	rd1 R1/L1-=490 bp, r2 R2/L1 470 bp.

**SET12B POL specific for *EGHV1,2,3,4,5* (second step)**

All use rd1 R1/L1, rd2 R2/L1, can add rd3 R2/L2 and R2/L3

**EGHV1**A/B R1-1	5’- CTYTACTGACTCCTGAAGTACTCC-3’

EGHV1A/B R1-2	5’- GYCATCCAGGCCTCAAACCGGGTG-3’

EGHV1A/B L1-2	5’-GTCAAGGCCTGCACCGATGCTGGTG-3’

EGHV1A/B L1-1	5’-CCTAAAGTGGGCTTCGGGATCTCC-3’

**EGHV2** R2-1	5’-CCCTCATAACATCAAAGGAGCTAC-3’

EGHV2 R2-2	5’-CACCCAGAACTAAAGGCGGAGGTG-3’

EGHV2 L2-1	5’-CTTTGATAAGGGATAAGGGATTACAG-3’

**EGHV3**A/B R3-1 5’-CCCAYGAGAAACTACAYATGC-3’

EGHV3A/B R3-2 5’-GCATGGCAATCTAARRCCTGAG-3’

EGHV3A/B L3-1 5’-CTGAATCTGGCATCCGGATCATGTTG-3’

**EGHV4** R4-1	5’-CCCTAATTTCCCACGAGAAACTG-3’	EGHV4 R4-2	5’-CATGCATGGTGACCTAAGGCCCG-3’

EGHV4 L4-1	5’-GACCCTAAATCTTGCATTTGGCTC-3’

EGHV4 NEW L4-2	5’-CTGAGGGCGTCTGGGGTAATTGACTC-3’

EGHV4 NEW L4-3	5’-CACAGGGAAGAACACCAGATGCCACACC-3’

**EGHV5**A/B R5-1	5’-CCCTCATTCCACACAATGAGCTTCATTTGC-3’

EGHV5A/B R5-2	5’-CATTTGCATTCTCATCTTACACCAAACG- EGHV5A/B L5-1	5’-GGGACGACGCAAGGTACGGTATAACCC-3’

Rd2A,2B,3A,3B (multiple alternative combinations)

**SET 15/16A U71/gM specific for *EEHV3/4/7***

R1 LGH6793B	5’-GCCATCGGGATCCGGAAAARACGCC-3’

R2 LGH9209	5’-CCACCGATACTACATGGGAAACATAGCC-3’ (same as LGH9137)

L4 LGH9211	5'-GGCAACATCACCGTAATGTACGTGGTTTGG-3’

L2 LGH9210	5’CCAGTACGATAAGATCTACCTGGACGA-3’ (same as LGH9138) L1 LGH6792B	5’-GGGCATCTTCTTCACCCTCATAGCCATC-3’

rd1 R1/L1 = 700-bp, rd2 R1/L2 = 670-bp; rd3 R2/L4 PCR = 640-bp.

**SET 16D U71/gM specific for *EEHV3B* and *EEHV7A/B***

R3Bsp	LGH9682	5’-GCCTTCGAGGAGCTATCGGATGAAGAGA-3’

L3Bsp	LGH9684	5’-GAGGTACAAACCCTTGGTGGTGCTCACA-3’

R3COM LGH9685 5’CTRCTGGACGAC GAAGCCTTCGAGGAGC3’

R7COM LGH9686 5’-CTGTTGGACGACGACGCGTTCGAGGACC-3’

L3+7COM LGH9687 5’-GCGTTGCCTCMGGATTAYTAYCACAAC-3’	Lalt3COM LGH9210 =(9138) 5’-CCAGTACGATAAGATCTACCTGGACGA-3’	rd1 R3/L3+7 =560-bp, rd2 R3B/L3+7 =545-bp, R3B/L3B =510-bp. **SET 18C U73(OBP) specific for GC-rich *EEHV3/4/7***

R1 LGH9279 5’-CGAGCCCCTATGGGGGTCTGGNAAGAC-

R2 LGH9280 5’-GACSGCGGCYATGATCTCGTGGCTCA- L4	LGH9283 5’-GGCACGAGGCGACGCACGCCAGGAACTTG- L0	LGH9288	5’-GACTGGTATACTGACATCATGTCGGGACC-3’

rd1 R1/L0 =940-bp, rd2R1/L4 =360-bp, rd3 R2/L4 = 345-bp

**SET 26 U48.5/U49 (gH-TK)**

R1 LGH9985 5’CTGTTCCTCTGTCTCCACGCGCCTCAGCGT-3”

R2 LGH9986 5'-AGCATGGAGTACAACGGCATGTAGCTGA-3'

L2 LGH9987 5'-GTATAGGTGTAGGTAAGACGTCACTGTTC-3’

L1 LGH9988 5'-CTCACCGTTTACCTAGAAGGTTGTATAGGTG-3'

rd1 R1/L1 = 360-bp, rd2A R1/L2 = 300-bp, rd2B R2/L1 = 280-bp, rd3AB R2/L2= 250-bp

**SET 26D new alternative specific for PAN *EEHV3/4/7* GC-branch**

R1	LGH11242: 5’-CCTGGCGAAAGGACCAGAGCATGGAG-3’

R2	LGH11243: 5’-GCAGGTACTGGTGCAGGGGAAACAC-3’

L2	LGH11244: 5’-CTGGACCCAGTGGTTTCCCGAAAACG-3’

L1	LGH11245: 5’-CGTCACTGTTCAAATACGCCGCGGATAAC-3’

rd1 R1/L1= 375-bp, rd2A R2/L1 = 320-bp, rd 2B R1/L2 = 305bp, rd 3AB R2/L2 = 250-bp **SET 27 U81(UDG)**

R1 LGH9989 5‘-AGGTAT TGGTTGGCTTTCACGAAGTG-3‘

R2 LGH9990 5‘-CTGGCGGCCGCCAGGGGGGAAGGGTGAGC-3‘

L2 LGH9992	5’-GCCGACGGCCTGGCCTTCTCCACGGGTGACGG-3’

L0 LGH11149 5‘-CGAAGAGTGGGTCGCCTTTCTAGA-3‘

rd 1R1/L0 = 520-bp, rd 2A R1/L2 = 435-bp, rd2B R2/L0= 470-bp, rd R2/L2 = 360-bp;

**SET 28 E54(vOX2-1) (*EEHV1A/1B/6* only)** R1 5’ LGH837 5’- ATGCTTCAGAGAAAGTAAGGTAC-3’.

R2alt LGH132 5’-GGTCGTAACGCAAGATGAGCGAG-3’	L1 LGH8472 5’-GTGTTGCCGCCACGATGCTTCTACG-3’	rd1 R1/L1 = 910-bp; rd 2 R2/L1 = 740-bp

**SET 29A U51(vGPCR1) specific for *EEHV3/7***

R1 LGH10784 5’-CGTTCCCGCTGTGGATCTACCAGGC-3’

R2 LGH10785 5’-CAGCGGGATGTTCTACACCATG-3’

R3 LGH10786 5’-GTCACCTTCGATCGCTGGTACTG-3’

L2 LGH10787 5’-GCTCTGTCCCGAGTACAGGTAGCAG-3’ L1 LGH10788 5’-CATGCCGTAGAACGGCCCTTGGG-3’

rd1 R1/L1 = 490bp, rd2A R1/L2 = 310bp, , rd2B R2/L1 = 420, rd3AB R3/L2 =<400bp

**SET 30 E4(vGCNT1) specific for *EEHV4/7***

LGH10789 vGCNT1 EEHV7R1	5’-AGGGCCATCTACGCCCCGCAGAACCT-3’

LGH10790 vGCNT1 EEHV7 R2 5’-TCCTGGAGTCGCGTGCAGGCCGA-3’

LGH10791 vGCNT1 EEHV7 L2	5’-CATGTCCGAGGTGTCGTACTTG-3’

LGH10792 vGCNT1 EEHV7 L1	5’-TCGTAGTAGTGCCACTTGACGAACCTG-3’

rd1 R1/L1 = 660-bp rd2A R2/L1 = 515bp, rd2B R1/L2 = 525bp, rd3AB R2/L2 =415-bp

**SET 35 U14 specific for *EEHV3***

LGHU14-R1	5’-GTCTTCGCAARCCGCGCTGGTCGGC-3’

LGHU14-R2	5’-GTCTCTGGATGTTCTGGGCATCCAG-3’

LGHU14-L2	5’-CCGTGTATGAARTACGTRTGTAAC-3’	LGHU14-L1	5’-CGAGCCGATATCGTCGACGATCTG-3’	rd1 R1/L1 = 920-bp, rd2A R1/L2 =870-bp, rd2B R2/L1 = 860-bp, rd3AB R2/L2 = 810-bp

**SET 36 U42(MTA) specific for *EEHV3A/B* includes across ex1/ex2 boundary**

LGHU42-R1	5’-GTAGTCGAAGACAGCGCGTTCTAG-3’

LGHU42-R2	5’-GACAACATGTTCGACACTCCGGTC-3’

LGHU42-L2	5’-GACGTTCGGCAGCTTTCCCGTGTA-3’

LGHU42-L1	5’-GACGAACTTTTTCTCCATGAGCATGG-3’	rd1 R1/L1 = 900bp, rd2A R1/L2 = 860bp, rd2B R1/L2 = 870- bp= rd3AB R2/L2 = 830-bp

**SET 37 U43(PRI) specific for *EEHV3A/B***

LGHU43 R1	5’-CAGGTGCTCTTCGCGACCGAATAC-3’

LGHU43 R2	5’-TGCGGATGCGATGTTGACCACCT-3’

LGHU43 L2	5’-CAGAACGCTATGATCGTTTCGTC- 3’	LGHU43 L1	5’-CTGACGCGGGTAGAGTTTTCGCACG-3’	rd1 R1/L1 = 760bp, rd2A R1/L2 = 720bp, rd2B R2/L1 = 710bp, rd3AB R2/L2 = 670-bp

**SET 38 U82(gL)-E37(ORF-Oex3) specific for *EEHV3/4/7***

LGH11558 New R1	5’-CGTTCACGGGCGGYGGGTASACGATC-3’

LGH11559 New R2	5’-GGGTASACCGATCTCGGAGACCCG-3’

LGH11560 New L5A 5’-GACGCGTTTATCGTCACRGGGAGGG-3’

LGH11561 New L4	5’-TTYGACACTCCCGCCCTCCTTGACG-3’

LGH11562 New R3	5’-CACYCTYAGTTCCTCSGTRTTATA C-3’

LGH11563 New R4	5’-GAA TAGAYTTAACRCCGGTAGGAAC-3’

LGH11564 NewL3	5’-GTTCCTACCGGYGGTGTTAAATCTATTC-3’

LGH11565 New L2	5’-CAGATGATAACGTCCCTSGACAAGTG-3’

LGH11566 New L1	5’-CAGATYACGTATGTYTTTRATCCG-3

RdA: rd1 R1/L4 =680-bp, rd2A R1/L5 =640-bp, rd2B R2/L4 =66-bp, rd3A R2/L5 =620-bp; Rd2B: rd1 R3/L1=705bp, rd2A R3/L2 =610-bp, rd2B R4/L1 =565-bp, rd3A R3/L3 =535-bp, rd3B R4/L2 =565-bp. Multiple combinations over 1800-bp.
